# Frontotemporal dementia-like disease progression elicited by seeded aggregation and spread of FUS

**DOI:** 10.1101/2024.06.03.593639

**Authors:** Sonia Vazquez-Sanchez, Britt Tilkin, Fatima Gasset-Rosa, Sitao Zhang, Diana Piol, Melissa McAlonis-Downes, Jonathan Artates, Noe Govea-Perez, Yana Verresen, Lin Guo, Don W. Cleveland, James Shorter, Sandrine Da Cruz

## Abstract

RNA binding proteins have emerged as central players in the mechanisms of many neurodegenerative diseases. In particular, a proteinopathy of <u>fu</u>sed in <u>s</u>arcoma (FUS) is present in some instances of familial Amyotrophic lateral sclerosis (ALS) and about 10% of sporadic FTLD. Here we establish that focal injection of sonicated human FUS fibrils into brains of mice in which ALS-linked mutant or wild-type human FUS replaces endogenous mouse FUS is sufficient to induce focal cytoplasmic mislocalization and aggregation of mutant and wild-type FUS which with time spreads to distal regions of the brain. Human FUS fibril-induced FUS aggregation in the mouse brain of humanized FUS mice is accelerated by an ALS-causing FUS mutant relative to wild-type human FUS. Injection of sonicated human FUS fibrils does not induce FUS aggregation and subsequent spreading after injection into naïve mouse brains containing only mouse FUS, indicating a species barrier to human FUS aggregation and its prion-like spread. Fibril-induced human FUS aggregates recapitulate pathological features of FTLD including increased detergent insolubility of FUS and TAF15 and amyloid-like, cytoplasmic deposits of FUS that accumulate ubiquitin and p62, but not TDP-43. Finally, injection of sonicated FUS fibrils is shown to exacerbate age-dependent cognitive and behavioral deficits from mutant human FUS expression. Thus, focal seeded aggregation of FUS and further propagation through prion-like spread elicits FUS-proteinopathy and FTLD-like disease progression.

## Background

Fused in sarcoma (FUS) is an RNA binding protein that normally localizes predominantly in the nucleus, however it mislocalizes and aggregates in the cytoplasm in some instances of familiar amyotrophic lateral sclerosis (ALS) and in 10% of frontotemporal lobar degeneration (FTLD), one of the most frequent forms of early-onset dementia (1-3). The 526-amino-acid FUS protein includes a C-terminal non-classical PY nuclear localization signal (NLS) which contains most of the ALS-linked mutations and a N-terminal low complexity, glycine-rich, prion-like domain (4). Although an important difference from transmissible spongiform encephalopathies (TSEs) is that prions behave like infectious agents, prion-like diseases belong to a group of protein misfolding neurodegenerative diseases that are characterized by the abnormal aggregation of defined host proteins (e.g., Amyloid β (Aβ) and tau in Alzheimer’s disease, α-synuclein in Parkinson’s disease, mutant polyglutamine repeats in Huntington’s disease, and TDP-43 in ALS and FTLD). Aβ, tau, α–synuclein, and TDP-43 inclusions have been shown to develop in a stereotypical, age-dependent manner in particular brain regions from which they appear to spread (5-7).

Increasing evidence supports a model whereby misfolded proteins released from a cell harboring pathological inclusions act on recipient cells to form *de novo* pathology by corrupting endogenous normal proteins to adopt pathological conformations. Injection of sonicated fibrils from either disease-associated α-synuclein, tau, or Aβ peptides unilaterally into mouse brains expressing the respective mutant protein (8-10), induces the spread of aggregates far from the site of injection, accelerating disease and enhancing neuronal loss. The repetition of this process has been proposed to underlie cell-to-cell propagation of pathological proteins throughout the brain (11, 12). Accumulating evidence supports cell-to-cell templated propagation of Aβ, tau, α–synuclein, and huntingtin (13-18). For ALS, evidence from cell culture has suggested spread from cell-to-cell from the dipeptide repeat (DPR) proteins encoded by hexanucleotide expansion in *C9orf72* (19, 20) and seeding and spread of SOD1 have also been reported with mutant SOD1 transgenic mice (21-24). Additionally, TDP-43 aggregates from FTLD patients and recombinant TDP-43 preformed fibrils have been proposed to induce prion-like spread pathology of the protein both in cultured cells and transgenic mice expressing cytoplasmic TDP-43 (25-29).

Neuropathological evidence from a small number of patients is consistent with the hypothesis of FUS pathology spreading within the central nervous system (CNS), including *1)* clinical symptoms often start focally and spread as disease progresses (30-32) and *2)* FUS cytoplasmic inclusions have been observed in several regions of the CNS of ALS and FTLD-FUS patients with similar spatial patterns as in FTLD-Tau or FTLD-TDP-43 forms (2, 33-35). That said, FUS inclusions vary markedly, presenting distinct density and shapes between cases (3, 35-39). Initial *in vitro* evidence for FUS seeding potency was provided by Nomura et al. who described that FUS-LCD fibrils carrying the G156E mutation seed wild-type FUS *in vitro* and in cell culture (40). Here we show that focal injection of pre-assembled human FUS fibrils in adult mouse brains induces *de novo* aggregation of endogenous human ALS-associated mutant FUS or human wild-type FUS and seeding of a spreading pathology through the nervous system that initiates neurodegeneration and compromises cognition.

## Methods

### Animals

The generation of the humanized FUS animals was described before (41). All the mice used in this report were maintained on a pure C57BL/6 background. For this study, we used 16 months old males and females with the mFUS^KO^/hFUS^R521H or WT^ genotype. All experimental procedures were approved by the Institutional Animal Care and Use Committee of the University of California, San Diego.

### Protein purification

Protein purifications were performed as described before (42). Briefly, HA-tagged FUS^R495X^ expression construct was generated using a pGST-Duet construct which contains a TEV-cleavable site, resulting in a GST-TEV-HA-FUS^R495X^ protein (43). All proteins were expressed and purified from *E. coli* BL21 CodonPlus (DE3)-RIL cells under native conditions. Protein expression was induced adding 1 mM IPTG for 16h at 16°C. *E. coli* bacterial cells were lysed on ice by sonication in Phosphate-Buffered Saline (PBS) supplemented with protease inhibitors (cOmplete, EDTA-free, Roche Applied Science). The protein was purified over pre-packed Glutathione Sepharose High Performance resin column (GSTrap HP columns, Cytiva). One-step purification of glutathione S-Transferase (GST) tagged FUS protein was performed using Akta Pure fast protein liquid chromatography (FPLC) system (Cytiva) at 4°C. GST-HA-FUS^R495X^ protein was eluted in 50 mM Tris-HCl, pH 8, 200 mM Trehalose, and 20 mM L-glutathione reduced. His-SOD1 protein was purified over pre-packed Ni Sepharose High Performance HisTrap HP (GE) using an AKTA pure chromatography system at 4°C and eluted with 50 mM Tris pH 7.4, 100 mM NaCl and 400 mM Imidazole. The following Molecular Weight Markers were used: Carbonic Anhydrase from bovine erythrocytes (29 KDa, Sigma), Albumin, bovine serum (66KDa, Sigma) and b-Amylase from sweet potato, (200KDa, Sigma). Eluted proteins (GST-HA-FUS^R495X^, and His-SOD1) with the expected size were collected and concentrated to final concentration of 12 mM using Amico Ultra centrifugal filter units (10 kDa molecular weight cut-off; Millipore). All proteins after purification were centrifuged for 15 min at 14,000 rpm at 4°C to remove any aggregated material. Protein concentration was calculated by Coomassie Blue with BSA protein as standard, and by colorimetric Bradford assay (Bio-Rad). For protein storage at -80°C glycerol (30%) was added.

### Protein fibrilization

FUS fibrilization was induced as described by Gasset-Rosa et al. (42). GST-HA-FUS^R495X^ protein was thawed and buffer exchanged into FUS assembly buffer at 4°C (50 mM Tris-HCl, pH 8, 200 mM trehalose, 1 mM DTT, 20 mM glutathione). TEV protease was added to GST-TEV-HA-FUS^R495X^ (4 µM) in FUS assembly buffer for 3 hours to induce seed formation. Next, high salt storage buffer (40 mM HEPES pH7.4, 500 mM KCl, 20 mM MgCl2, 10% glycerol, 1 mM DTT) was added for 3 hours to separate the seeds (43). FUS fibrilization was initiated by adding 5% of FUS seeds to GST-TEV-HA-FUS^R495X^ (4 µM) and TEV protease in FUS assembly buffer for 24 hours at 22°C. His-SOD1 fibrilization was induced as described in (42, 43). Finally, fibrils were dialyzed using slide-A-Lyzer MINWE Dialysis Units (10 kDa molecular weight cut-off; Thermo Fisher Scientific) in PBS for 3 hours and sonicated at 45% 45s just before injecting them into the animals.

### Transmission Electron Microscope

300-mesh Formvar/carbon coated copper grids (Ted Pella) were glow-discharged and loaded with fibril protein samples (10 µl). Next, grids were stained with 2% (w/v) aqueous uranyl acetate (Ladd Research Industries, Williston, VT). Excessive liquid was removed and grids were air dried. Grids were examined using a Tecnawe G2 Spirit BioTWIN transmission electron microscope equipped with an Eagle 4k HS digital camera (FEI, Hilsboro, OR).

### Stereotactic injections

All surgical procedures were performed using aseptic techniques. Injections were performed using 33-gauge needles and a 10 µl Hamilton syringe (Hamilton, Switzerland). 16-month-old mFUS^KO^/hFUS^R521H^, mFUS^KO^/hFUS^WT^ or non-transgenic mice were injected with 10 µg of sonicated HA-FUS^R495X^ fibrils, 10 µg of sonicated SOD1 fibrils, 10 µg HA-FUS^R495X^ monomer or PBS as control, following the stereotactic coordinates using bregma as a reference: anteroposterior – 2.5 mm, mediolateral – 2.0 mm and dorsoventral at -1.8 mm (hippocampus) and -0.8 (cortex).

### Behavioral tests

For each behavioral assay, a cohort of n=5-6 animals per group for the non-transgenic genotype (FUS fibrils or PBS-injected) and n=11-12 animals per group for the mFUS^KO^/hFUS^R521H^ genotype (FUS fibrils or PBS-injected) was assessed where experimentalist was blinded to genotypes. No increased mortality was observed in any of the groups.

#### Open field test

The open field area consisted out of a square white Plexiglas (50 × 50 cm2) open field illuminated to 600 lx in the center and mice were placed in the center. The mice were allowed to explore the area for 10 mins. An overhead Noldus camera was used to monitor their movement with Ethovision XT software. Mice were tracked for multiple parameters, including distance traveled, velocity, center time, frequency in center as described in (44).

#### Rotarod

The rotarod test was performed as described in (45). A Rota-rod Series 8 apparatus (Ugo Basile) was used. Before the trial was initiated, the mice were placed on the stationary rotarod for 30s for training. Each mouse was given three trials per day, with a 60s inter-trial interval on the accelerating rotarod (4–40 r.p.m. over 5 min) for five consecutive days. The latencies to fall were automatically recorded by a computer.

#### Novel object recognition test

This behavioral assay was performed as described in (41). Mice were individually habituated to a 51cm x 51cm x 39cm open field for 5 min and then tested with two identical objects placed in the field. Each mouse was allowed to explore the objects for 5 min. After three such trials (each separated by 1 min in a holding cage), the mouse was tested in the object novelty recognition test in which a novel object replaced one of the familiar objects. Behavior was video recorded and then scored for contacts (touching with nose or nose pointing at object and within 0.5 cm of object). Habituation to the objects across the familiarization trials (decreased contacts) is an initial measure of learning and then renewed interest (increased contacts) in the new object indicated successful object memory. Recognition indexes were calculated using the following formula: # contacts during test/(# contacts in last familiarization trial + # contacts during test). Values greater than 0.5 indicate increased interest, whereas values less than 0.5 indicate decreased interest in the object during the test relative to the final familiarization trial.

### Immunofluorescence

Mice were intracardially perfused with 4% paraformaldehyde (PFA) in PBS and the full brain was post-fixed in the same 4% PFA for 2 hours and transferred to 30% sucrose in PBS for at least 2 days. Brain was embedded in HistoPrep (Fisher Chemical) and snap frozen in isopentane (2-methylbutane) cooled at – 40°C on dry ice. 35 µm brain cryosections were cut using a Leica 2800E Frigocut cryostat at -20°C and stored as free-floating sections in 1X PBS + 0.02% Sodium Azide at 4°C. The free-floating brain sections were washed 3 times, 10 min in 1X PBS and then incubated in blocking solution (0.5% Tween-20, 1.5% BSA in 1X PBS) for 1 hour at room temperature (RT) followed by overnight incubation at RT in antibody diluent (0.3% Triton X-100 in 1X PBS) containing the primary antibodies. The next day, sections were washed again 3 times, 10 min in 1X PBS and incubated with secondary antibody (Jackson Immunoresearch, diluted in 0.3% Triton X-100 in 1X PBS), washed again 3 times with 1X PBS and then incubated 10 min with DAPI diluted in 1X PBS (Thermo Fisher Scientific, 100 ng/ml). Sections were mounted on Fisherbrand Superfrost Plus Microscope Slides (Thermo Fisher Scientific) with Prolong Gold antifade reagent (Thermo Fisher Scientific). Full brain images were acquired with the Nanozoomer Slide Scanner (Hamamatsu©) and visualized in NDP.view2 software. Close-up images of brain sections displaying individual neurons were acquired with the FV1000 Spectral Confocal (Olympus) at 60X magnification or the spinning disk confocal Yokogawa X1 confocal scanhead mounted to a Nikon Ti2 microscope with a Plan apo lamda 100x oil NA 1.45 objective and Plan apo lamda 60x oil na 1.4 objective.

To quantify of FUS aggregates, brain coronal sections were carefully matched to compare similar anatomical regions, keeping track of the injected and non-injected side and immunostained with FUS, A11, OC and LOC antibodies as well DAPI. The percentage of DAPI positive cells with mislocalized aggregated FUS from similar area sizes within the cortex and hippocampus was counted.

To quantify the levels of nuclear and cytoplasmic FUS in neurons, brain coronal sections were matched to compare similar anatomical regions and immunostained with FUS, NeuN and DAPI. Images from the cortex were segmented using NeuN to identify neurons, as well as the outline of the Neuron itself. DAPI was used to segment the nucleus and record nuclear FUS intensity in neurons. Substracting the DAPI mask to the NeuN mask was used to define the cytoplasm of neurons and record FUS cytoplasmic intensity.

To quantify the number of neurons in the mouse brains, coronal brain OCT sections were immunostained with NeuN and nuclei were stained with DAPI. The hippocampal dentate gyrus region contained mostly NeuN-positive cells. For its quantitation, DAPI-positive cells were counted manually in 3–5 consecutive sections per animal using Fiji software. For the motor cortex region, NeuN-positive cells were counted to exclude glia cell nuclei. Careful matching of the sections to compare similar anatomical regions was performed for each set of mice.

### Antibody list

| Antibody | Supplier | Catalog # | Dilution |
| --- | --- | --- | --- |
| Anti-FUS | Bethyl | A303-839A | IF 1:500 |
| Anti-NeuN | Genetex | GTX133127 | IF 1:500 |
| Anti-p62 | Progen | GP62-C | IF 1:500 |
| Anti-TDP-43 | Proteintech | 10782-2-AP | IF 1:500 WB 1:1000 |
| Oligomer A11 | Invitrogen | AHB0052 | IF 1:500 |
| Anti-P-Ubiquitin (Pser65) | Millipore | MAB1510 | IF 1:500 |
| Anti-amyloid fibrils LOC | Millipore | AB2287 | IF 1:500 |
| Anti-Amyloid Fibrils OC | Millipore | AB2286 | IF 1:500 |
| Anti-Glial Fibrillary Acidic Protein (GFAP) | Millipore | MAB360 | IF 1:1000 |
| Anti-Hsp90 (C45G5) | Cell Signaling Technology | 4877S | WB 1:5000 |
| Anti-Iba1 | Wako | 019-19741 | IF 1:500 |
| Anti-FUS | Bethyl | A300-294A | WB 1:1000 |
| Anti-TAF15 | Bethyl | A300-309A | WB 1:1000 |
| Anti-GAPDH | Millipore | CB1001 | WB 1:5000 |

### Serial fractionation and Western Blot

Mouse brains were homogenized in high-salt (HS) buffer (4 ml/g; 50 mM Tris pH 7.5, 750 mM NaCl, 5 mM EDTA and protease and phosphatase inhibitor mix). Then the sample was diluted 1:10 with Pierce™ IP Lysis Buffer [25 mM HEPES (pH7.4), 150 mM NaCl, 1 mM EDTA, 1% NP40, 5% V/V glycerol, protease and phosphatase inhibitor, and 1/100 V/V benzonase (endonuclease)]. The sample was incubated on ice for 30 min, sonicated and centrifugated for 1hour at 10 000 x g at 4°C. The supernatant was used as the soluble fraction and the pellet was resuspended in Laemmli SDS-loading buffer and used as the insoluble fraction. 10% Bis-Tris gels were used for immunoblotting and equal volumes of samples were loaded. For antibodies, see antibody list.

### Serial fractionation and dot blot

Serial fractionation was performed as in (41). Mouse cortices were homogenized in high-salt (HS) buffer (4 ml/g; 50 mM Tris pH 7.5, 750 mM NaCl, 5 mM EDTA and protease inhibitor mix), then centrifuged for 30min at 45 000 x g at 4°C resulting in the HS fraction. Next, the pellet was homogenized in 500 ml of HS buffer + 1% Triton X-100 and 1M sucrose and centrifuged 30min, 4°C at 45 000 x g (HS + Tx fraction). Then the remaining pellet is suspended in urea buffer (2 ml/g; 7M urea, 2M thiourea, 4% CHAPS, 30 mM Tris pH 8.5), centrifuged at 45 000 x g and for the remaining pellet 2 ml/g of SDS loading buffer was added. Equal volumes were spotted onto a nitrocellulose membrane. For antibodies, see antibody list.

### Statistical analysis

Statistical analysis was performed using GraphPad Prism. All data is shown as mean ± standard error of the mean (SEM). The Kolmogorov-Smirnov normality test was used to evaluate the distribution of the data. If comparing two normal distributed groups, t-test was used. In case of comparing more than two normally distributed groups, data were compared by one-way analysis of variance (ANOVA) with Dunnets post-hoc tests. When data were not normality distributed and homoscedastic, the Kruskal-Wallis test was used with Dunn’s multiple test as post-hoc. When P-values were lower than 0.05, significance was noted in the figure as: *P<0.05, **P<0.01, ***P<0.001, ****P<0.0001. Detailed information is shown in each figure legend.

## Results

### Amyloid-like fibrils of FUS induce aggregation and time-dependent spread of human mutant FUS^R521H^

FUS pathology is present in rare sporadic ALS and familial ALS (46), but is a hallmark of nearly 10% of the sporadic FTLD patients, known as FTLD-FUS (37). FUS aggregation is almost universally found in sporadic FTLD-FUS patients with inclusions that are tau- and TDP-43-negative (38, 46, 47). While we (41) and others (48-51) demonstrated that FUS aggregation is not required for disease initiation in mice expressing ALS-linked FUS mutations, but rather for its misaccumulation in axons and cytoplasm, respectively, here we devised to test whether FUS aggregation contributes to disease progression. To do this, we exploited our humanized FUS mice in which mouse FUS is replaced by the human full-length FUS gene encoding either wild-type FUS (mFUS^KO^/hFUS^WT^) or ALS-linked FUS^R521H^ (mFUS^KO^/hFUS^R521H^), the latter of which develops late onset progressive motor and cognitive deficits without detectable cytoplasmic FUS aggregation (41). We expressed and purified full-length recombinant human FUS^R495X^ protein HA-tagged on its amino terminus (Fig.1A), incubated with FUS^R495X^ seeds at 22°C for 24 hours to generate spontaneously assembled, amyloid-like fibrils *in vitro* (Fig. 1B). We selected FUS^R495X^ fibrils as the initial seeds as they would be predicted to evade rapid disaggregation by endogenous Karyopherin-β2 since FUS^R495X^ lacks the PY-NLS region recognized by Karyopherin-β2 (52) and allow to distinguish between the endogenous FUS and the exogenous fibrils by immunostaining using an antibody against the 500-526 amino acid peptide sequence of FUS protein that is missing in the FUS^R495X^ fibrils. Those fibrils were then sonicated (Fig. 1B) and injected unilaterally after disease initiation into the cortex and hippocampus of 16-month-old humanized FUS mice (mFUS^KO^/hFUS^R521H^, Fig. 1C).

**Figure 1:**
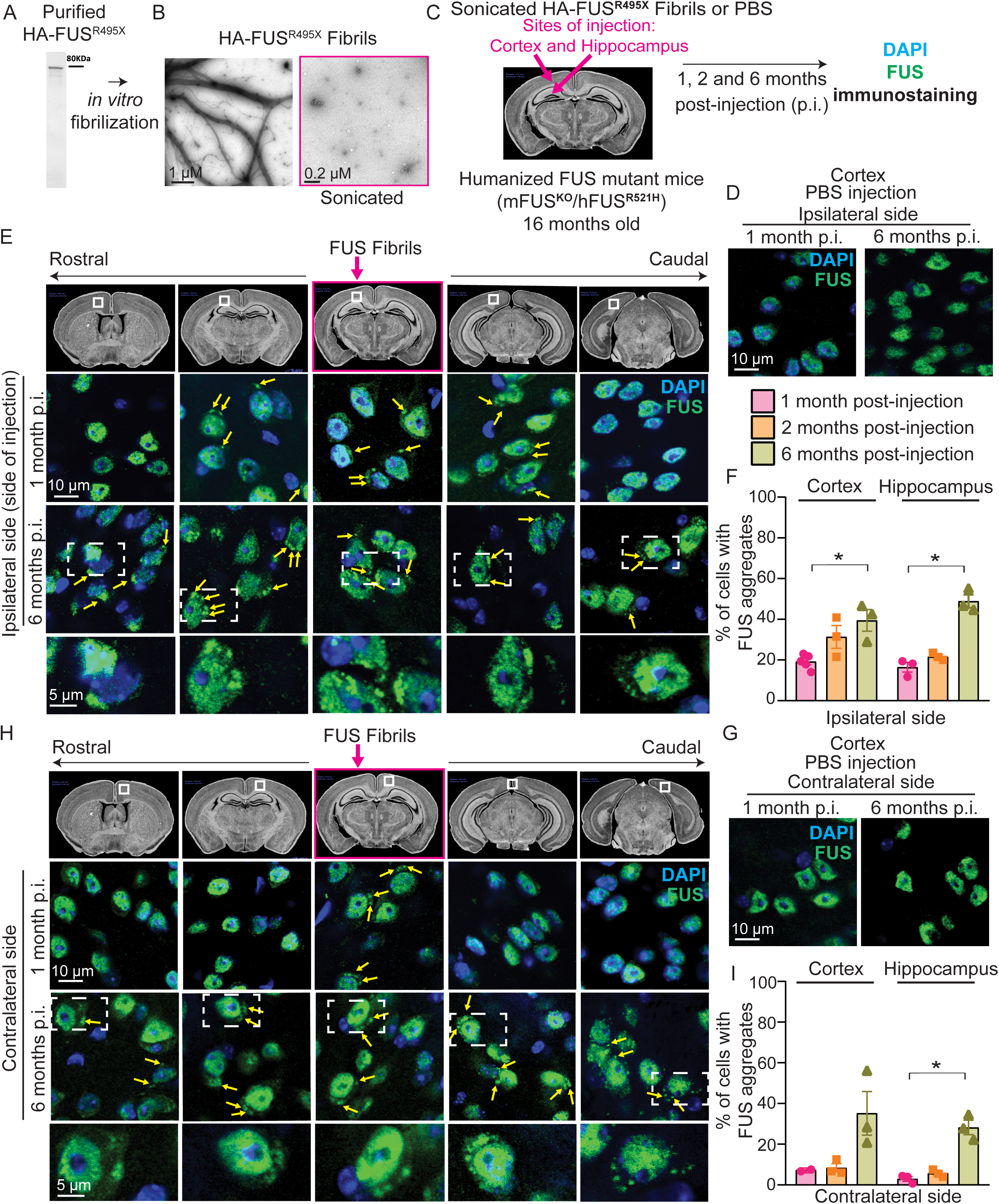
HA-FUS^R495X^ fibrils induce human FUS mislocalization and aggregation in aged humanized mutantFUS (mFUS^KO^/hFUS^R521H^) mice. **A.** Coomassie blue staining of recombinant HA-FUS protein. **B.** Electron micrograph of fibrils of HA-FUS^R495X^ recombinant protein purified from bacteria (Left panel). HA-FUS^R495X^ fibrils after sonication before inoculating them into mice (Right panel). Scale bars: 1 µm (before sonication), 0.2 µm (after sonication). **C.** Sonicated HA-tagged FUS^R495X^ fibrils were injected unilaterally into the cortex and hippocampus of 16 months old humanized, mutant FUS mice (mFUS^KO^/hFUS^R521H^). **D,E.** Immunostaining of FUS (green) and DAPI (blue) of the side of the mouse brain hemisphere (ipsilateral side) injected either with PBS (D) or with HA-FUS^R495X^ fibrils (E) after 1 month and 6 months post-injection. Scale bars: 10 µm, inset: 5 µm. The top panel illustrates the regions of the brain that were analyzed and the site of injection (pink box). Yellow arrows indicate cytoplasmic FUS aggregates at 1- and 6-months post-injection (p.i.). **F.** Quantification of the percentage of cells containing endogenous cytoplasmic FUS aggregates in the cortex and hippocampus at the injection (ipsilateral) side 1-, 2- and 6-months post-injection. N=3 animals. Kruskal-Wallis test with Dunn’s multiple test post-hoc p-values: cortex p = 0.0429 and hippocampus p = 0.0219. Data is presented as mean ± SEM. **G,H.** Immunostaining of FUS (green) and DAPI (blue) of the opposite side of the mouse hemisphere (contralateral side) that was injected either with PBS (G) or with HA-FUS^R495X^ fibrils (H) after 1 month and 6 months post-injection. Yellow arrows indicate FUS cytoplasmic inclusions after 1- and 6-months post-injection. Scale bars: 10 µm, inset: 5 µm. **I.** Quantification of the percentage of cells containing endogenous FUS aggregates in the cortex and hippocampus at the contralateral side over 1-, 2- and 6-months post-fibril injection. N=3 animals. Kruskal-Wallis test with Dunn’s multiple test post-hoc p-values: p = 0.0225.

The fate of the sonicated fibrils was followed over time using immunodetection of the HA-epitope tag to mark the pre-formed FUS fibrils. Three hour-post-injection exogenously produced HA-tagged FUS fibrils were immunodetected using the HA-epitope tag against the pre-formed FUS fibrils and were found to focally distribute into the cortex and hippocampus within a 150 µm-area anterior-posterior from the injected coordinates (Fig. S1A, left panel and Fig. S1B). Fibrils persisted for the following 3 days and were predominantly immunodetected in the cytoplasm, suggesting their uptake from the cells (Fig. S1A, middle panel). However, one week after injection no pre-formed FUS fibrils were detected, consistent with clearance of the sonicated FUS fibrils (Fig. S1A, right panel). Focal injection provoked activation of microglia and astrocytes in hippocampus and cortex (Fig. S2) at 3 days post-injection on the site of injection, which was absent in the contralateral side. This astrocytic/microglial activation was only transient as it was not detected at 3 hours and was mitigated by 1-week post-injection (Fig. S2). Since astrocytes and microglia can internalize and degrade added external aggregates *in vitro* (53, 54), glia cells may contribute to the clearance of the injected fibril material.

To test whether focal injection of sonicated FUS fibrils into cortex and hippocampus of humanized mutant FUS^R521H^ mice recapitulates FUS pathology and if so whether it propagates with time beyond the injection site, we further analyzed brain sections one, two, and six months post-injection (Fig. 1C). FUS protein partially redistributed to the cytoplasm and was recruited into ∼1-5 FUS immunopositive inclusions, while FUS remained almost entirely in the nucleus in PBS-injected mice (Fig. 1C-E). FUS cytoplasmic inclusions were observed in brain regions that were in contact with exogenous FUS seeds, but also in adjacent regions without any apparent contact with the injected amyloid-like fibrils, including the contralateral side of the cortex and hippocampus which also exhibited cytoplasmic inclusions of FUS (Fig. 1D-I). FUS aggregates progressively spread into superficial layers found beyond the brain areas that were directly connected to the cortex and hippocampus regions of the injection site (Fig. 1E,H).

Over time, FUS aggregation was immunodetected throughout the whole hemisphere at the level of injection and in wider areas of the opposite hemisphere (Fig. S3), indicating a time-dependent spread of FUS aggregates to distal regions rostrally and caudally from the focal injection of FUS fibrils. At the site of injection (ipsilateral side), 19, 31, and 39% of the cortical and 16, 22, and 49% of hippocampal cells harbored FUS inclusions at 1-, 2-, and 6-months post-injection, respectively, *versus* 3, 7, and >30% on the contralateral side at both brain regions (Fig. 1F,I) while none were observed in PBS injected mice (Fig. 1D,G). Overall, these data demonstrate that *1)* exogenous FUS fibrils seed *de novo* aggregation of endogenous human FUS^R521H^ in a spatial-, temporal-dependent manner and *2)* FUS pathology spreads to distal sites, including within the non-injected hemisphere (albeit cells within the contralateral side of the injected hemisphere exhibited a reduced number of FUS inclusions relative to the site of injection).

### Neither monomers of FUS nor sonicated fibrils of SOD1 produce cytoplasmic aggregation of human mutant FUS^R521H^

In contrast to cytoplasmic aggregate induction and spreading of endogenously expressed FUS when mice were unilaterally injected with sonicated FUS fibrils in the cortex and hippocampus, FUS remained almost exclusively nuclear without detectable cytoplasmic aggregates in animals injected with FUS monomers (Fig. 2A,B). Additionally, we generated fibrils (Fig. 2C-E) of recombinant wild-type superoxide dismutase (SOD1) (43) and focal injection of the sonicated SOD1 fibrils into mFUS^KO^/FUS^R521H^ brains did not provoke aggregation or mislocalization of FUS locally or distally to the injection sites (Fig. 2E). Therefore, the recruitment of endogenous mutant FUS to sonicated fibril-induced FUS mislocalization and aggregation was unique to the injection of sonicated FUS fibrils.

**Figure 2:**
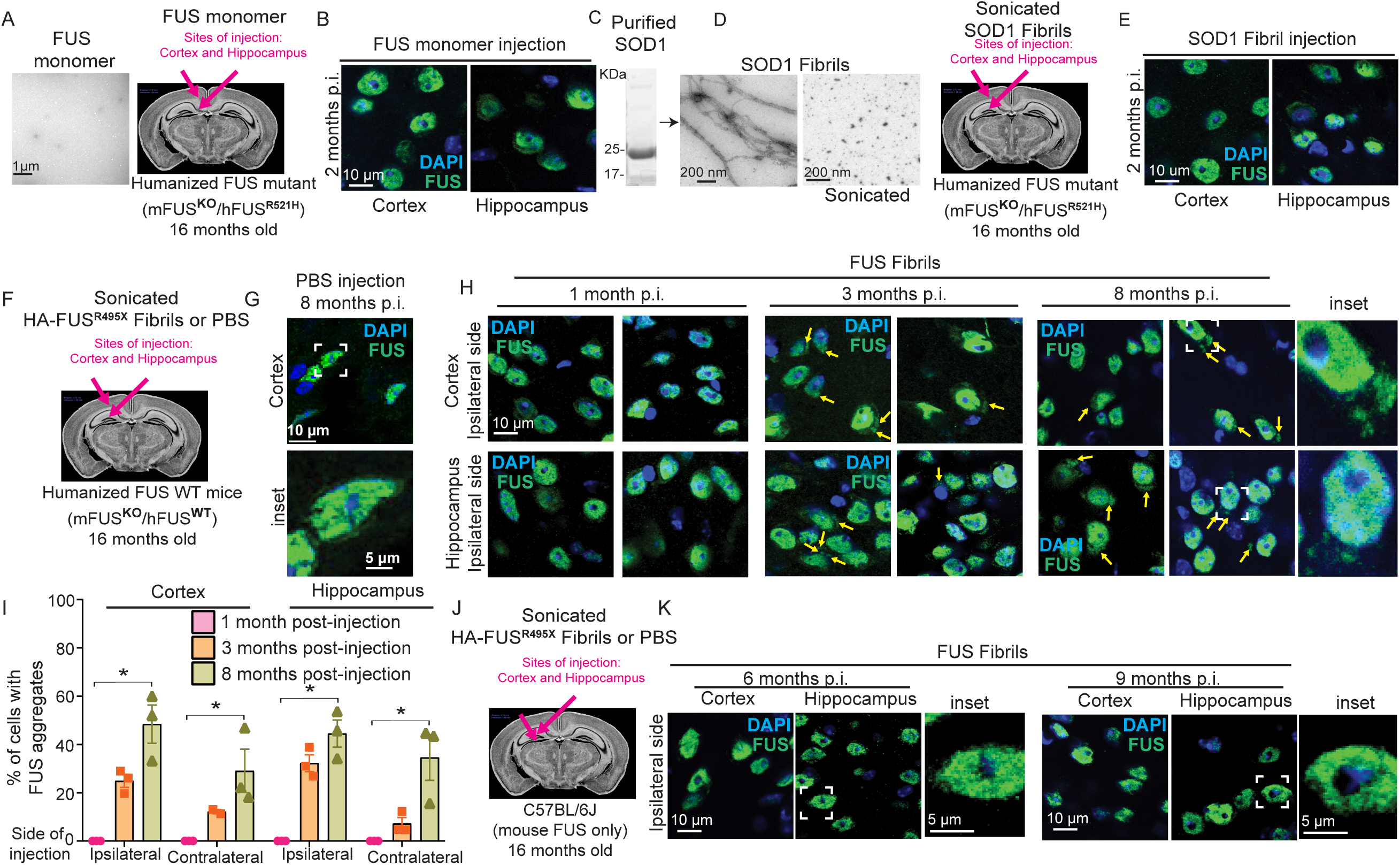
HA-FUS^R495X^ fibrils induce aggregation and spreading of human FUS WT but not of mouse FUS. **A.** Electron micrograph of FUS monomeric protein (scale bar: 1 µm) which was injected into 16-month old mFUS^KO^/hFUS^R521H^ mice. **B.** Immunostaining of FUS (green) and DAPI (blue) of the side of the mouse brain in which FUS monomers were injected after 2 months post-injection. Scale bars: 10 µm. **C**. Coomassie blue staining of recombinant His-SOD1 protein. **D.** Electron micrograph of His-SOD1 fibrils obtained from recombinant protein purified from bacteria (left panel) and sonicated His-SOD1 fibrils before inoculating them into 16 months old humanized, mutant FUS mice (mFUS^KO^/hFUS^R521H^). Scale bar: 200 nm. **E.** Immunostaining of FUS (green) and DAPI (blue) of the side of the mouse brain in which FUS monomers were injected after 2 months post-injection. Scale bars: 10 µm. **F**. Sonicated HA-tagged FUS^R495X^ fibrils were injected unilaterally into the cortex and hippocampus of 16 months old humanized, FUS wild-type mice (mFUS^KO^/hFUS^WT^). **G**. Immunostaining of a PBS-injected mFUS^KO^/hFUS^WT^ mouse brain, 8 months post-injection using a FUS (green) antibody. **H**. Immunostaining of FUS (green) and DAPI (blue) of the side of the mouse brain hemisphere (ipsilateral side) injected with HA-FUS^R495X^ fibrils. Yellow arrows indicate FUS cytoplasmic aggregates after 1-, 3- and 8-months post-injection. Scale bars: 10 µm, inset: 5 µm. **I.** Quantification of the percentage of cells with endogenous human FUS aggregates in the cortex and hippocampus at the ipsilateral and contralateral side, 1-, 3- and 8-months post-injection. N=3 animals. Kruskal-Wallis test with Dunn’s multiple test post-hoc p-values: cortex ipsilateral p = 0.0190, cortex contralateral p = 0.0312, hippocampus ipsilateral p = 0.0299 and hippocampus contralateral p = 0.0190. Data is presented as mean ± SEM. **J,K.** Immunostaining of FUS (green) and DAPI (blue) of the side of injection (ipsilateral side) in non-transgenic C57BL/6J mice (mouse FUS) brains injected with HA-FUS^R495X^ after 6- and 9-months post-injection. Scale bars: 10 µm, inset: 5 µm.

### Mutant FUS accelerates FUS aggregation induced by injected FUS fibrils

To test if human wild-type FUS can be seeded to aggregate (as is seen in examples of sporadic ALS and FTLD (46)), FUS^R495X^ fibrils were focally injected at single sites within the cortex or hippocampus of 16-month-old humanized mFUS^KO^/FUS^WT^ mice in which both endogenous mouse FUS alleles had been inactivated (Fig. 2F). While *de novo* aggregation of endogenous human wild-type FUS was induced to a level comparable to that generated in humanized mutant FUS^R521H^ mice similarly injected, aggregation was accelerated by two months in the mutant FUS animals (Fig. 1 and 2F-I). Specifically, after injection of FUS^R495X^ fibrils, aggregation of endogenous human wild-type FUS was not observed until 3 months, while aggregation of endogenous mutant FUS was observed the first month post-injection. By 3 months post-injection, cytoplasmic wild-type FUS aggregates were found in 25% and 32%, respectively, of cells in the ipsilateral cortex and hippocampus, with spreading producing aggregates in 12% and 7%, respectively, of cells in the contralateral hemisphere (Fig. 2I). By eight months post-injection, wild-type FUS-containing aggregates were immunodetected throughout the brain in areas outside the injection site and the percentage of cells with cytoplasmic FUS aggregates rose to 46% in the ipsilateral side (in cortex and hippocampus), and 20% in cortex and 32% in hippocampus of the contralateral side (Fig. 2I). Overall, these findings support that *1)* focally injected FUS^R495X^ fibrils seed aggregation of wild-type endogenous human FUS and *2)* the induced wild-type FUS-containing inclusions propagate beyond the injection site to the opposite hemisphere, with the kinetics of spreading slower than for mutant FUS (Fig. 1 and 2F-I).

### A species barrier to FUS aggregate seeding

Sequence variations between species have been well established to create a species barrier for prion seeding and spread (55). To test for the presence of a similar species barrier for FUS seeding, human FUS^R495X^ fibrils were assembled, sonicated, and injected unilaterally into cortex and hippocampus of C57BL/6J mice exclusively expressing mouse FUS (Fig. 2J). Examinations at timepoints up to 9 months post-injection revealed that mouse FUS continued to be almost exclusively nuclear (Fig. 2J,K), with no aggregates detectable at any time point, consistent with an interspecies transmission barrier that limits the capability of sonicated human FUS fibrils to seed aggregation of mouse FUS.

### Injected FUS fibrils increase insolubility of endogenous FUS

To determine whether cytoplasmic FUS inclusions revealed by immunostaining acquire the characteristics of FUS inclusions found in postmortem patient material, we used a combination of immunocytochemistry and biochemistry at multiple time points post-fibril injection (Fig. 3A). Within 1 month post-fibril injection aggregated endogenous FUS (Fig. 3B) acquired pre-amyloid properties as determined by immunodetection with the A11 antibody that has been established to recognize a peptide backbone epitope common to pre-amyloid oligomers (56). By 6 months post-injection, an overwhelming majority of FUS aggregates (79.7%± 5.4) were A11-positive (Fig. S5), while as expected no such signals were present either in age-matched PBS control injected mice or within 3 hours after FUS fibril injection (Fig. 3B,C). Injection-induced human FUS inclusions in brains of FUS humanized mice were also immunopositive (Fig. S5) using antibodies previously reported to recognize mature, *in vivo* β-amyloid structures (51). Similarly, fibril-induced FUS aggregates co-localized with p62 and ubiquitin (as described in human FUS proteinopathies (2, 35, 57, 58)), but did not contain detectable levels of TDP-43 (Fig. 3D,E).

**Figure 3:**
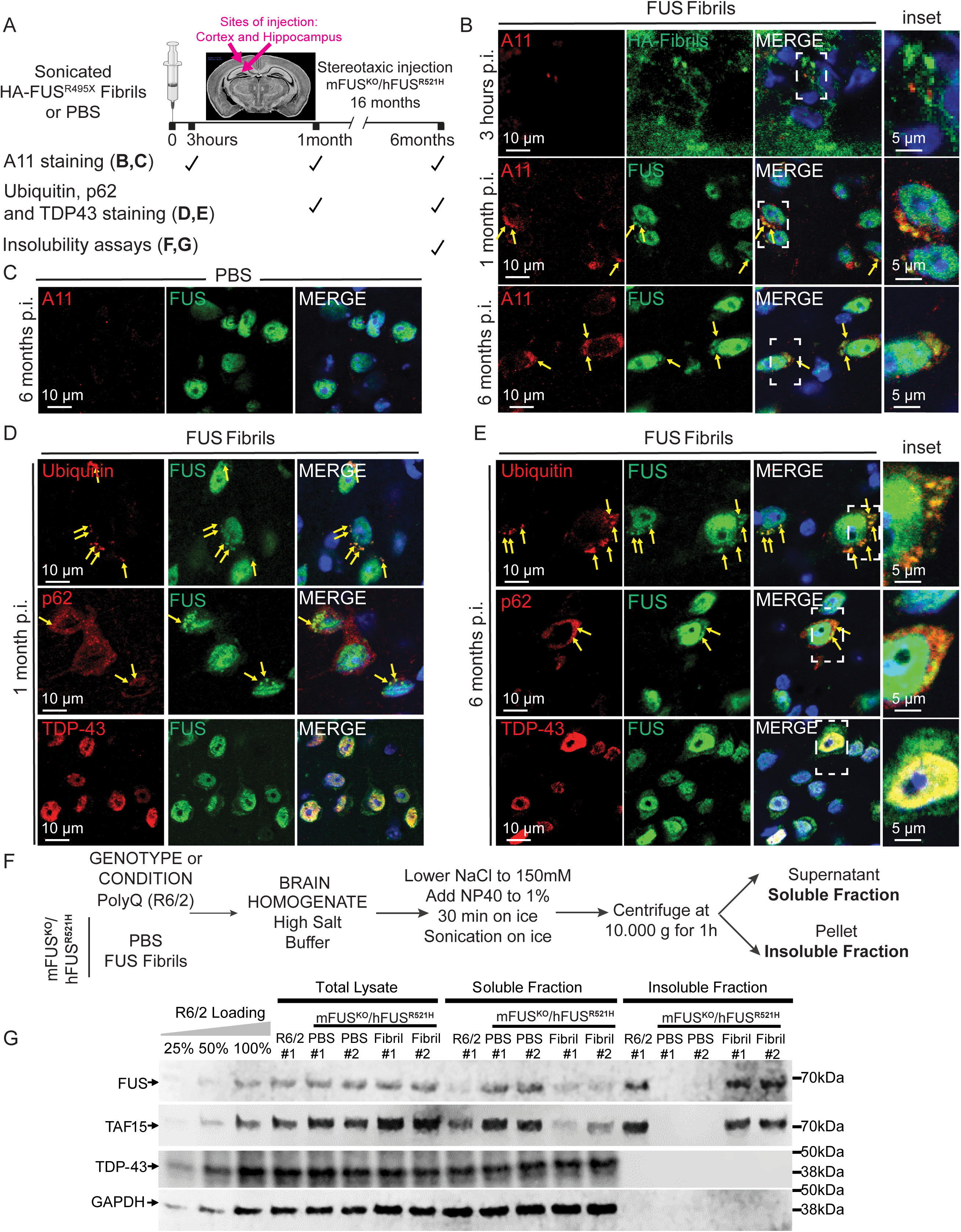
Human FUS aggregates are insoluble, display pre-amyloid properties and recapitulate features of human FUS pathology. **A.** Schematic overview of the timepoints at which FUS aggregation was analyzed using immunofluorescence-based assays and biochemical insolubility assays at 3 hours, 1 months and 6 months post-injection. **B,C.** Representative confocal images of mFUS^KO^/hFUS^R521H^ mouse brains injected either with HA-FUS^R495X^ fibrils at 3 hours, 1 months and 6 months post-injection (B) or with PBS at 6 months post-injection (C) immunolabelled using antibodies against the pre-amyloid oligomer marker A11 (red), HA after 3 hours post-injection (green) and FUS after 1 month and 6 months post-injection (green). Yellow arrows indicate co-localization between A11 and FUS cytoplasmic inclusions detected in fibril-injected mice. DAPI (blue) as nuclear counterstaining. Scale bars: 10 µm, inset: 5 µm. **D,E.** Representative confocal micrographs of mFUS^KO^/hFUS^R521H^ mouse brains injected with HA-FUS^R495X^ fibrils using ubiquitin/p62/TDP-43 (red) and FUS (green) antibodies after 1 month (D) and 6 months (E) post-injection. Yellow arrows indicate co-localization between either ubiquitin/p62/TDP-43 and FUS cytoplasmic aggregates. Scale bar: 10 µm, inset: 5 µm. **F**. Experimental outline of the serial fractionation of brain homogenates derived from mFUS^KO^/hFUS^R521H^ mice either PBS- or HA-FUS^R495X^ fibril-injected and R6/2 Huntington’s model mice as a positive control for FUS insolubility (59). **G.** Immunoblotting of the sequential biochemical fractions from mouse brains using anti-FUS, anti-TAF15 and anti-TDP-43 antibodies. Anti-GAPDH was used as loading control.

Analysis of brain homogenates from FUS-injected mice showed a marked increase in detergent-insoluble FUS compared with PBS-injected mice (Fig. 3F,G and Fig. S5E,F).

While in humanized FUS mutant mice (mFUS^KO^/hFUS^R521H^) most FUS remained soluble as we previously reported (41), an increase (compared to PBS injected brains) in insoluble FUS was detected in extracts from brains that were FUS fibril-injected (Fig. 3F,G). The FUS homologue TATA-binding protein-associated factor 15 (TAF15) (also known as TATA-binding protein-associated factor 2N), but not the RNA binding protein TDP-43, was also present in a detergent insoluble fraction, consistent with increased FUS and TAF15 insolubility reported in ALS/FTLD-FUS patients (57, 59) and extraction and imaging of filaments of TAF15 from such patient samples (59).

After 6 months of sonicated FUS fibril injection, the vast majority (90.9%±4.2) of cytoplasmic human endogenous FUS inclusions was present within neurons, with the remaining 10% in glia (Fig. S6A,B). Although the overall levels of nuclear and cytoplasmic FUS did not change 6 months post-injection of HA-FUS^R495X^ fibrils (Fig. S6C-E), the neurons bearing FUS cytoplasmic inclusions displayed a trend towards a decrease in the FUS nuclear/cytoplasmic ratio (Fig. S6F-H). Furthermore, formation of cytoplasmic FUS inclusions was accompanied by increased astrocytosis and microgliosis in fibril-injected mice (as revealed by increased immunoreactivity with GFAP and IBA1 antibodies, respectively) at 6 months, but not at 2 months post-injection (Fig. S7).

### Seeded aggregation of FUS provokes neurodegeneration

Single-dose injections of sonicated FUS fibrils (HA-FUS^R495X^) or PBS in the cortex and hippocampus were administered to cohorts of either non-transgenic mice or humanized FUS mutant mice (mFUS^KO^/hFUS^R521H^) and their behaviors were monitored using novel object recognition, rotarod and open field assays at timepoints prior to injection, and at 2 months and 6 months post-injection (Fig. 4A-B, Fig. S7). Cognitive impairments associated with mutant FUS expression (as we previously reported (41)) were significantly aggravated 6 months post-injection (Fig. 4A). While no exacerbation of disease was observed in humanized mutant FUS mice within the first two months after fibril-injection, rotarod performance was modestly decreased in fibril-injected humanized mutant mice 6 months post-injection (Fig. S8). In an open field assay, focal injection of mutant FUS fibrils also induced age-dependent deficits in the humanized FUS mice (Fig. 4B). Moreover, analysis of cortical and hippocampal sections revealed a significant neuronal loss in both hippocampus and cortex after FUS fibril injection compared to their PBS and non-injected controls (Fig. 4C-E), indicating exacerbation of behavioral deficits and neurodegeneration that correlate with seeded, prion-like spread of aggregated FUS.

**Figure 4:**
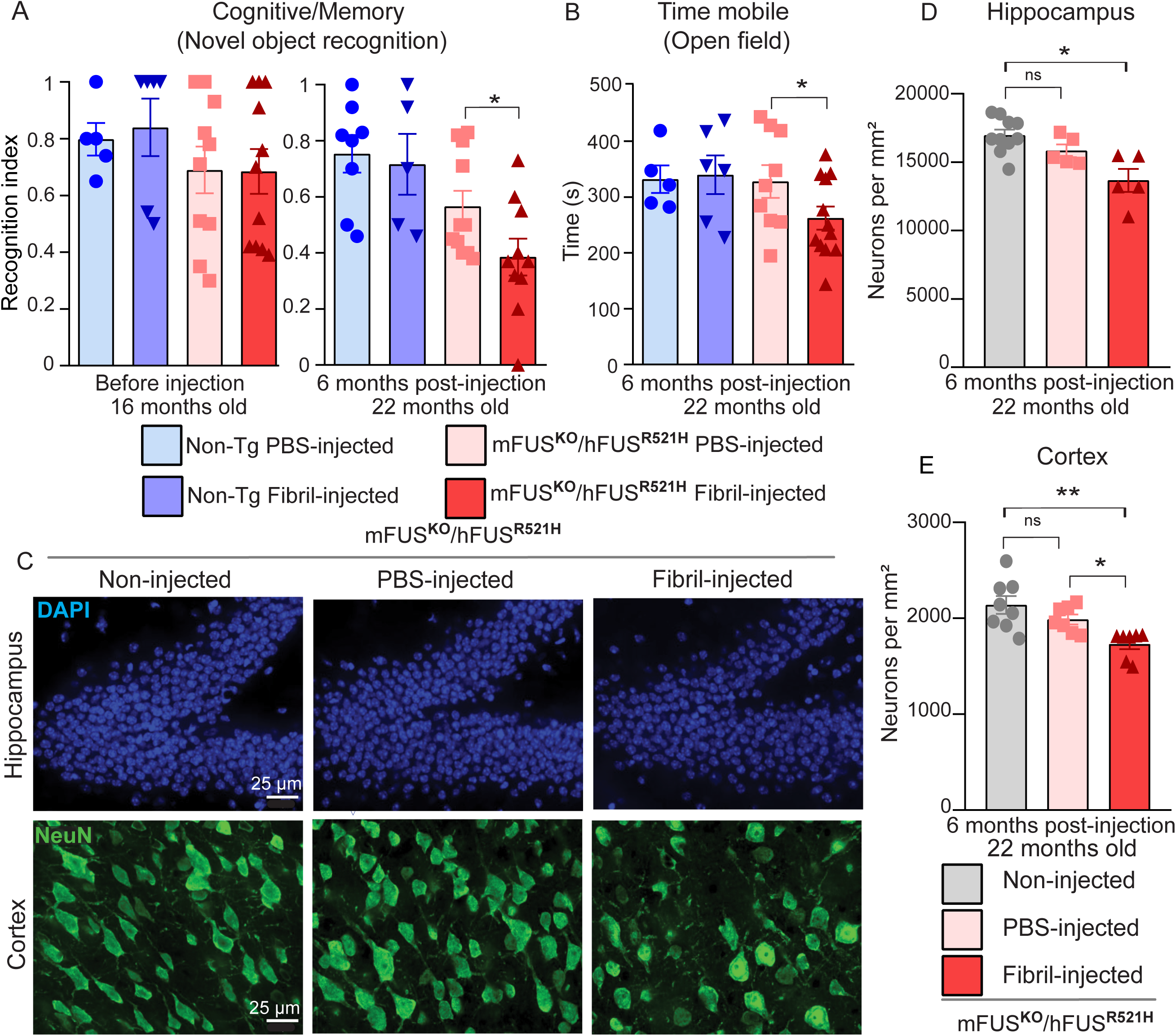
Human FUS aggregates exacerbate cognitive impairments and provoke behavioral deficits and neurodegeneration in ALS-FUS mice. **A.** Novel object recognition test was performed in 16 months (before injection) and 22 months old HA-FUS^R495X^ fibril-injected mFUS^KO^/hFUS^R521H^ animals (6 months post-injection) compared to PBS-injected controls and non-transgenic HA-FUS^R495X^ fibrils or PBS injected controls. N=5–12 animals per group. Unpaired t-test p-value = 0.0293. Data is presented as mean ± SEM. **B.** Open field test was performed in 22 months old HA-FUS^R495X^ fibril injected mFUS^KO^/hFUS^R521H^ animals (6 months post-injection) compared to PBS-injected controls and non-transgenic HA-FUS^R495X^ fibrils or PBS injected controls. N=5–12 animals per group. Unpaired t-test p-value = 0.0498. Data is presented as mean ± SEM. **C.** Representative immunofluorescence labelling for the neuronal marker NeuN (green) and DAPI in the hippocampus (upper panel) and cortex (lower panel) of humanized mutant mice mFUS^KO^/hFUS^R521H^ non-injected, and injected either with PBS or HA-FUS^R495X^ fibrils. Scale bar: 25 µm. **D,E.** Quantification of neurons in hippocampus (D) and cortex (E) in HA-FUS^R495X^ fibril-injected mice compared to PBS and non-injected controls. N=4 animals per condition. Data is presented as mean ± SEM. Kruskal-Wallis test with Dunn’s multiple test post-hoc p-values: hippocampus (D) p* = 0.0112 and cortex (E) p* = 0.0455 and p** = 0.0053.

## Discussion

We have developed a model of FTLD disease in mice through the dissemination of FUS pathology within the brain, affecting cognitive and motor functions. Focal injections of human FUS aggregates in the brain of humanized FUS mice induce *de novo* pathology of endogenous mutant or wild-type human FUS spreading within the brain in a spatio-temporal manner consistent with a model of transmission of pathology (in a prion-like fashion) throughout the brain. Endogenous FUS aggregation was observed at 1 month post-injection around the site of injection in the hippocampus region, a region associated with pathology in both ALS and FTLD patients (2, 60) and in the cortical region, where basophilic and FUS positive inclusions are found in neurons and glia cells of ALS cases, and are numerous in the middle and deep layers of the neocortex in FTLD (46, 61). Pathology after injection was found as well in brain areas distant from the injection sites, including in the contralateral hemisphere where FUS cytoplasmic aggregates were observed in a pattern mirroring the injected hemisphere similar as α-synuclein, tau and TDP-43 spreading in mice (9, 29, 62, 63). Our data contribute to the mounting evidence that prion-like transmission of misfolded proteins represents a common process in the pathogenesis of several neurodegenerative diseases, including α-synuclein in Parkinson’s disease (9, 17, 64), Aβ and Tau in AD (18, 65, 66), and SOD1 or TDP-43 in ALS (21, 25, 26, 28).

Injection of sonicated fibrils of hFUS^R495X^ induced aggregation and time-dependent spread of endogenous human mutant and wild-type FUS albeit the spreading mechanism remains unknown. FUS pathology may spread through adjacent cell-to-cell seed transfer, through anatomical neuronal connections, possibly in a diffusion-like manner, or (most likely) a combination of several mechanisms. The fact that FUS aggregates are found in the contralateral hemisphere suggests spread between hemispheres through brain commissures possibly the corpus callosum, but also through smaller anterior, posterior and hippocampal commissures (67). Injecting sonicated FUS fibrils in an area with known distal projections, such as the lateral geniculate nucleus, and evaluating the presence of FUS cytoplasmic aggregation in the synaptically connected visual cortex (68), would be valuable to determine if neuronal connectivity facilitates the spreading of FUS cytoplasmic aggregates. The time-dependent increasing accumulation of FUS cytoplasmic aggregates in the more distant brain areas after FUS-fibril injection further points to prion-like spreading mechanisms in the formation of FUS pathology. One cannot completely exclude that the injected sonicated FUS fibrils spread in a diffusion-like manner and seeded neurons in distal regions from the injection site, where they were undetectable right after injection. However, after the initial seeding event, a plausible route for FUS aggregates to appear in distal regions with time is spreading of human endogenous FUS seeds through the extracellular space via its release from dying cells, and/or through secretion (freely or through extracellular vesicles) and uptake into recipient cells, again via endocytosis and endosomal membrane rupture as it has been proposed for spreading of other prion-like proteins such as tau (69-71).

Great heterogeneity in the morphology of FUS cytoplasmic inclusions has been reported in human disease, at least some of which has been correlated with disease severity and FUS mutation in ALS cases (36). Moreover, FTLD-FUS pathology is divided into 3 different groups based on the morphology of the cytoplasmic inclusions and their deposition pattern (37). We observed round shaped cytoplasmic inclusions of FUS mostly in neurons that were ubiquitinated, p62 positive and TDP-43 negative. Moreover, cytoplasmic FUS aggregates and non-pathogenic nuclear FUS are detected in the same cell as reported before (61, 72). After 6 months of fibril injection, human mutant FUS further display enhanced insolubility together with enhanced TAF15 insolubility but not TDP-43. Both FUS and TAF15 were also detected in the detergent-insoluble fraction of the Huntington’s disease mice R6/2, indicating that TAF15 insolubility seems a secondary effect of FUS aggregation and is not due to exposure to injected FUS fibrils. However, it is possible that focal injection of amyloid-like FUS fibrils caused endogenous mouse TAF15 to form fibrils which also spread throughout the brain. It would thus be of interest to test if TAF15 depletion can prevent the spreading of cytoplasmic FUS aggregation in mice and whether injection of recombinant TAF15 fibrils can induce endogenous human FUS cytoplasmic aggregation Altogether, our model recapitulates the FUS and TAF15 shift in solubility and immunoreactivity for ubiquitin and p62 positive (but TDP-43 negative) that has been reported in human FTLD-FUS (3, 35-39, 59, 73).

Proteins with prion-like domains form pathological inclusions in many neurodegenerative diseases. The core region of the low complexity domain (LCD) of FUS is essential to form parallel β-sheet structures reminiscent of the amyloid-like proteins (74). Seeding is an important feature of amyloid-like aggregates, in which a piece of protein fibril can function as a structural template for facilitating the fibrillation of soluble protein molecules (75). Fibril-induced FUS cytoplasmic inclusions exhibit enhanced insolubility, supporting the idea that FUS inclusions could effectively transform soluble FUS into insoluble aggregates, resulting in the progressive dysfunction of FUS and cytotoxicity. Indeed, a seeded fibrillation of proteins and their intercellular transmission have been increasingly noticed as a molecular pathomechanism that describes the progression of several neurodegenerative diseases (76, 77).

Permissive prion transmission frequently depends on overcoming a species barrier, which is determined by a range of possible conformers of a particular prion, its sequence, as well as its interaction with cellular co-factors (78, 79). Here, we show the existence of a seeding barrier between human and mouse FUS. One such endogenous **‘**barrier**’** relevant to FUS proteinopathy may be the sequence differences between mouse and human FUS (which differ in 26 out of 526 amino acids), fifteen of which are located in the G-rich, prion-like domain believed to be a major factor in driving aggregation (Fig. S8) (80). Another plausible explanation is that mouse FUS is intrinsically less aggregation prone and cannot be seeded and/or spread. A future experiment that would decipher between these possibilities would be the injection of recombinant mouse FUS fibrils in non-transgenic wild-type mice to test if this will induce endogenous murine FUS aggregation and spreading, further supporting the idea of a species barrier.

FTLD patients with FUS inclusions only rarely harbour genetic alterations in FUS (81) and the majority of cases are sporadic (14). Most of the ALS-linked FUS mutations reported to date are localized in the NLS domain resulting in impaired nuclear transport of FUS and chaperoning by Karyopherin-β2. This is consistent with the finding that mutant FUS^R521H^ exhibit accelerated initial seeding and aggregation capacity compared to wild-type FUS, due to a reduced transport efficiency to the nucleus and increased retention in the cytoplasm.

Seeded aggregation of FUS provoked neurodegeneration and impaired mouse behaviour but the underlying molecular mechanisms mediating cell toxicity remain to be elucidated. ALS-linked mutations in FUS induce a gain of toxicity that includes stress-mediated suppression in intra-axonal translation, and synaptic dysfunction (41). With FUS fibril-injection we observed a portion of FUS mislocalized to the cytoplasm, clustering in visible inclusions that are widespread within the brain. Deletion of the NES in FUS strongly suppressed toxicity of mutant FUS in *Drosophila* (82), suggesting that the cytoplasmic localization of mutant FUS confers toxicity which is supported by the neurodegeneration we observed in both the hippocampus and cortex of fibril injected mice. In parallel, we observed gliosis at late time points of FUS pathology that might contribute to the damage of the tissue and translated in behaviour deficits since inflammation and astrocyte-mediated toxicity have been identified as part of the pathogenic process of ALS/FTLD (83). A natural follow up of this work is to characterize the composition of the cytoplasmic FUS inclusions and define the spatial transcriptomic changes provoked by FUS aggregation at different time points and brain regions as this may provide insights into the pathways of seeding, spreading and vulnerability or resistance. Deciphering such molecular mechanisms that underlie the spreading of FUS proteinopathy may offer avenues for therapeutic interventions by blocking spreading and thereby disease progression.

## Conclusion

Here we show that single focal injection of sonicated human FUS fibrils into aged brains of humanized FUS mice (in which ALS-linked mutant or wild-type human FUS replaces endogenous mouse FUS) induces FUS cytoplasmic aggregation, which recapitulates features of human FUS inclusions found in ALS/FTLD patients. Importantly, spread of FUS aggregates is shown to exacerbate FTLD-like disease induced by a disease-causing mutation and ultimately initiates neurodegeneration, thus providing the first *in vivo* evidence of spreading of templated FUS aggregation in an adult central nervous system.

### Abbreviations

Aβ: Amyloid β
ALS: Amyotrophic lateral sclerosis
DPR: dipeptide repeat
FTLD: Frontotemporal Lobar Degeneration
FUS: Fused in sarcoma
LCD: low complexity domain
NLS: Nuclear Localization Signal
PBS: Phosphate-Buffered Saline
SOD1: superoxide dismutase
TAF15: FUS homologue TATA-binding protein-associated factor 15
TSEs: transmissible spongiform encephalopathies

## Declarations

### Ethics approval and consent to participate

All animal experimental procedures were approved by the Institutional Animal Care and Use Committee of the University of California, San Diego, USA.

### Consent for publication

Not applicable for this study.

## Data availability

All data generated or analyzed during this study are included in this published article and available from the corresponding author on reasonable request.

## Authors’ contributions

FGR and SDC conceptualized and designed the study. SVS, BT, FGR, SZ, DP, MMD, JA, YV and NGP performed experiments and analyzed the data. LG and JS provided key reagents. SVS, BT, FGR, DWC and SDC wrote the manuscript, which was reviewed by all authors.

## Acknowledgements

We thank J. Santini at the UCSD Microscopy Core, E. Griffis and P. Guo at the UCSD Nikon Imaging Center and Y. Jones at Electron Microscopy Core Facility of UCSD for assistance with imaging and image analysis. We thank Z. Melamed, C. Chillon-Marinas, R. Maimon, J. Lopez-Erauskin, M. Baughn and S. Lu for their helpful discussions.

---

**Figure S1:**
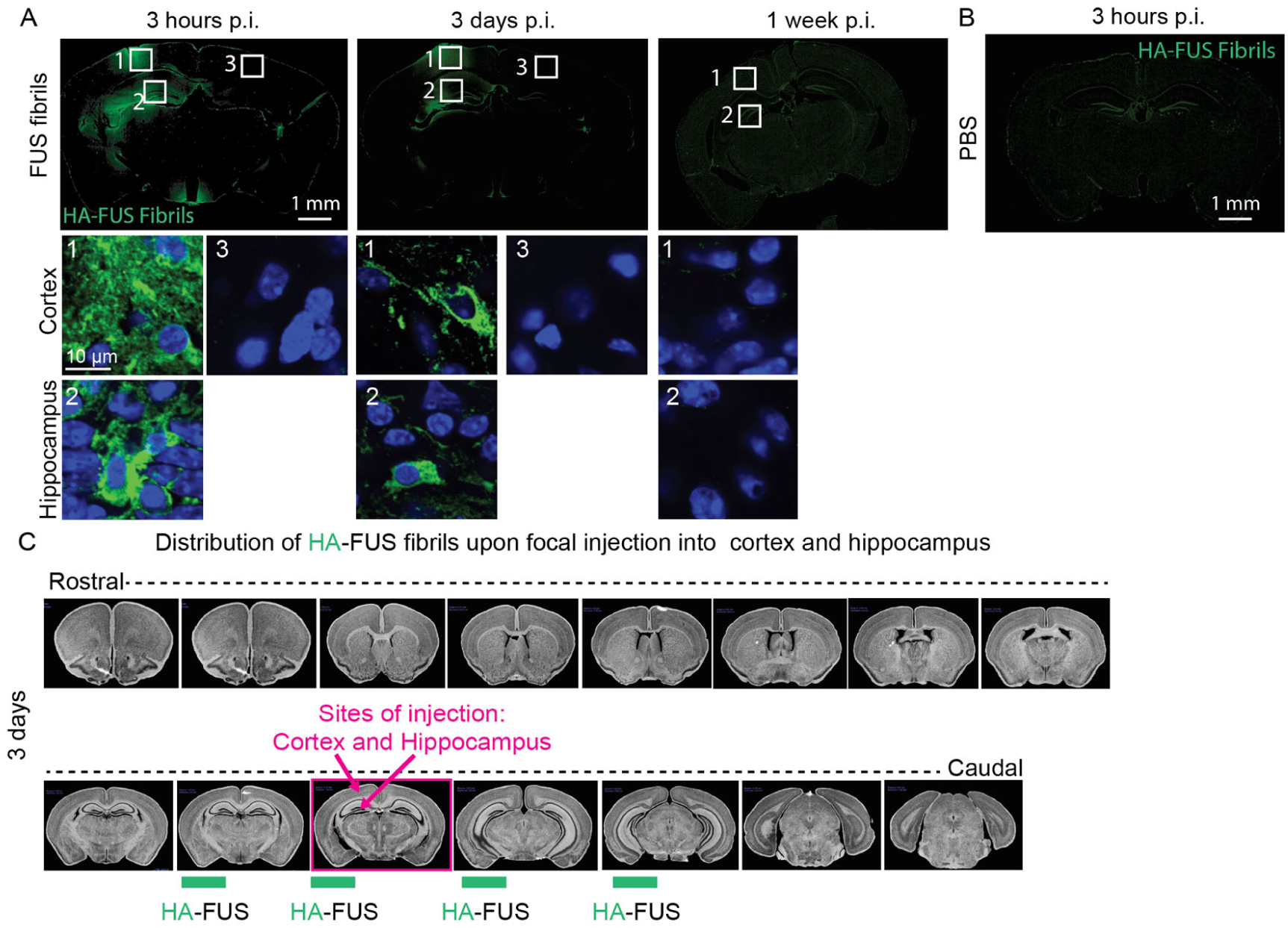
Distribution of focally-injected HA-tagged FUS^R495X^ fibrils in the mouse brain at 3 hours, 3 days and 1-week post-injection. **A.** Immunostaining of cortex and hippocampus of mFUS^KO^/hFUS^R521H^ mouse brain sections 3 hours, 3 days and 1-week post-injection with HA-FUS^R495X^ fibrils using an anti-HA antibody to detect the HA-tagged FUS^R495X^ fibrils. Number 1-2 correspond to the side of injection, number 3 corresponds with the contralateral side. DAPI as nuclear counterstaining. Scale bar: 10 µm. **B.** Immunostaining of mFUS^KO^/hFUS^R521H^ mouse brain sections 3 hours post-injection with PBS using an anti-HA antibody. **C.** Illustration summarizing the distribution of HA-FUS^R495X^ fibrils throughout the 3-day post-injection. Red dots indicate the sites where FUS aggregates are immunodetected.

**Figure S2:**
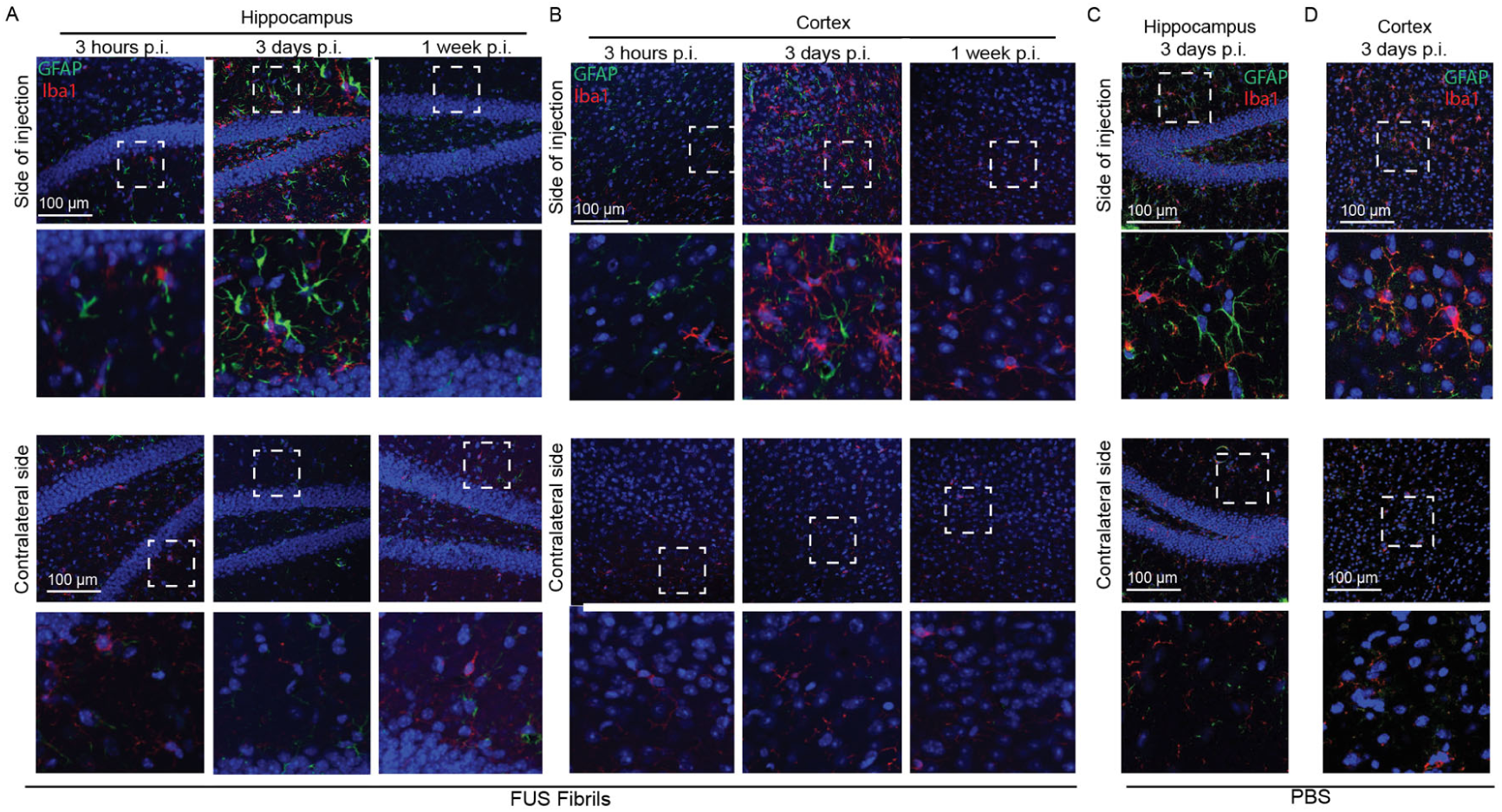
Focal stereotactic injections induce transient activation of glial cells. Representative confocal images of the site of injection in hippocampus (**A**) and cortex (**B**) of mFUS^KO^/hFUS^R521H^ mice injected with HA-FUS^R495X^ fibrils, 3 hours, 3 days or 1-week post-injection immunolabeled for GFAP (astrocytes) and Iba1 (microglia). DAPI as nuclear counterstaining. Representative confocal images of the site of injection in hippocampus (**C**) and cortex (**D**) of mFUS^KO^/hFUS^R521H^ mice injected with PBS, 3 days post-injection immunolabeled for GFAP (astrocytes) and Iba1 (microglia). DAPI is used as nuclear counterstaining. Scale bar: 100 µm.

**Figure S3:**
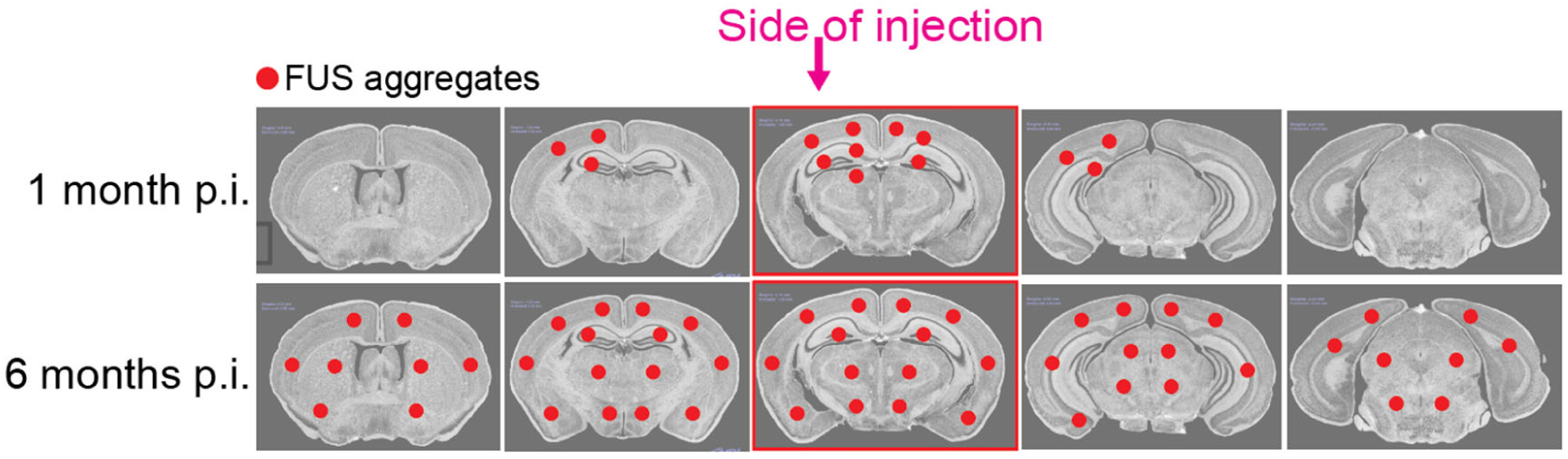
Distribution of FUS cytoplasmic aggregates in mFUS^KO^/hFUS^R521H^ mice induced by HA-FUS^R495X^ fibrils. Illustration summarizing the distribution of FUS cytoplasmic aggregates throughout the brain, 1- and 6-months post-injection. Red dots indicate the sites where FUS aggregates are immunodetected.

**Figure S4:**
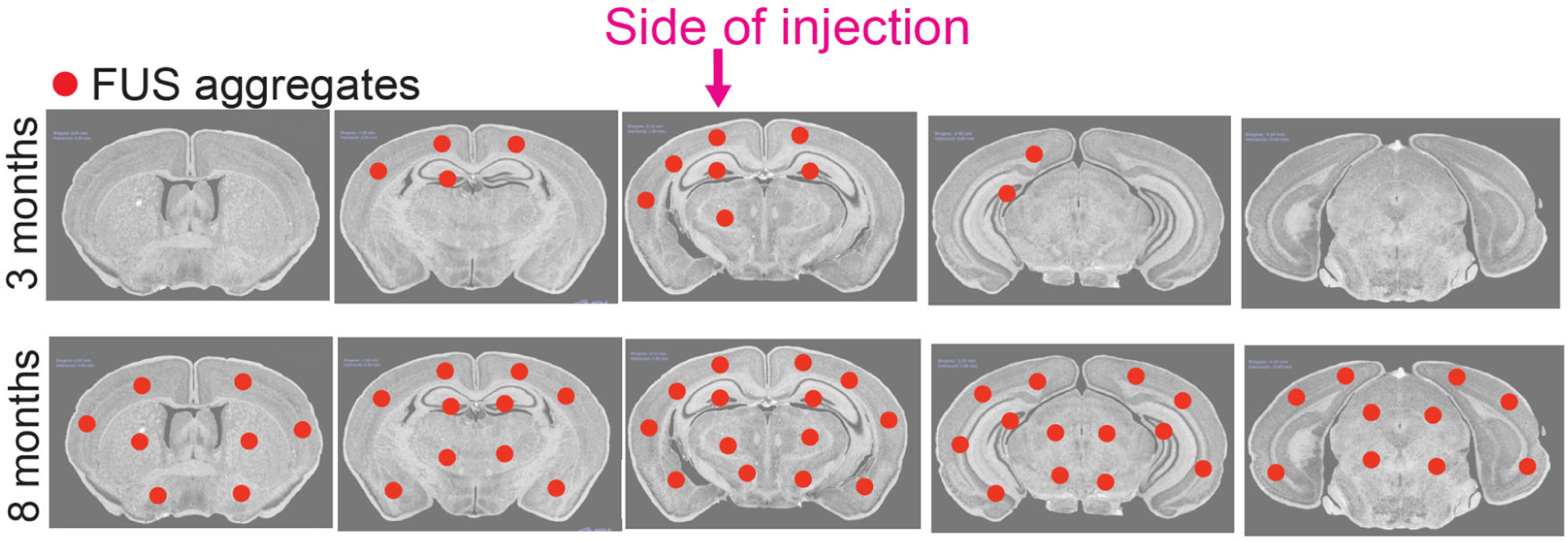
Distribution of FUS cytoplasmic aggregates in mFUS^KO^/hFUS^WT^ mice induced by HA-FUS^R495X^ fibrils. Illustration summarizing the distribution of FUS cytoplasmic aggregates throughout the brain, 3- and 8-months post-injection. Red dots indicate the sites where FUS aggregates are immunodetected.

**Figure S5:**
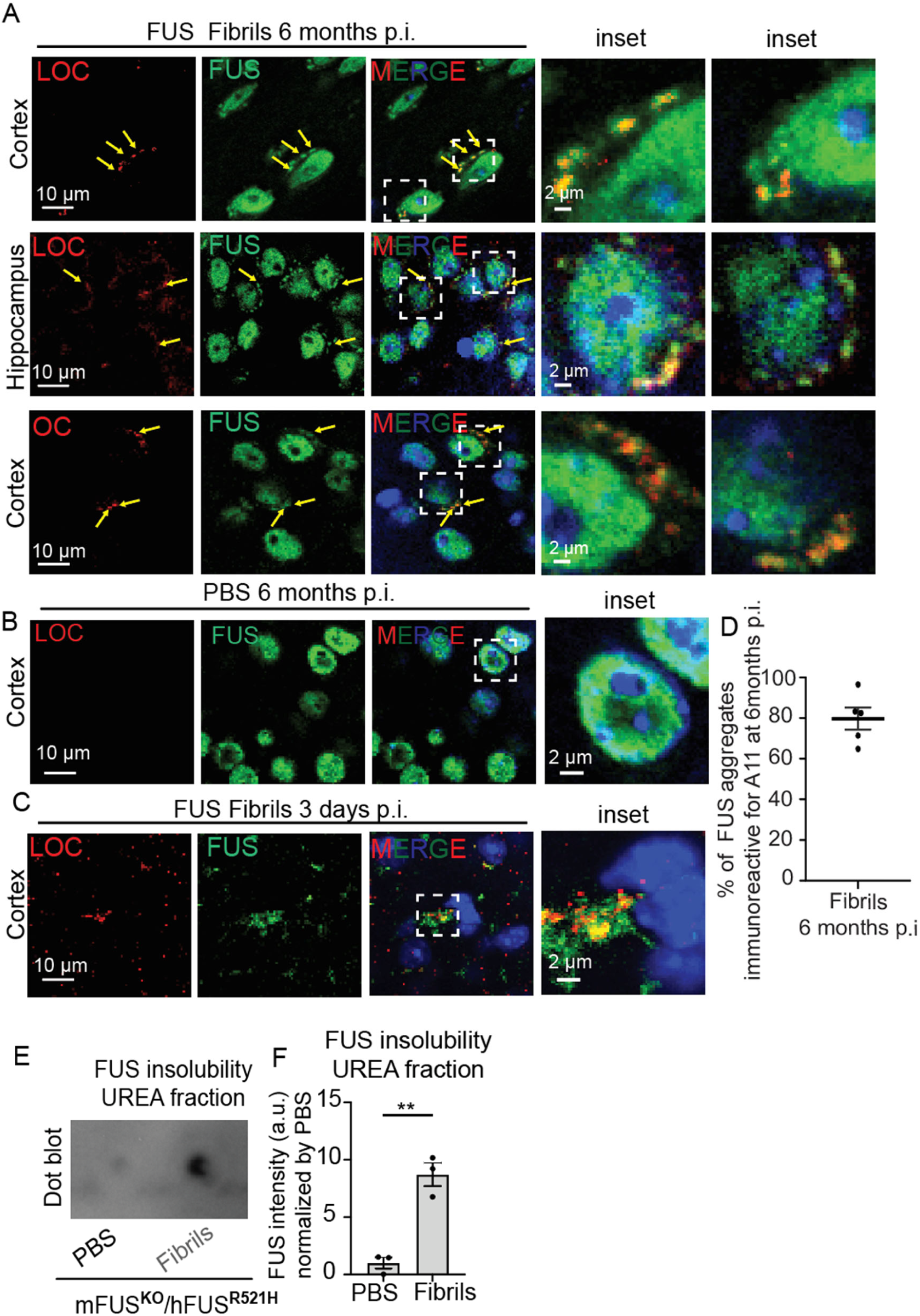
FUS aggregates display amyloid properties. **A,B.** Representative confocal micrographs of mFUS^KO^/hFUS^R521H^ mouse brains injected with HA-FUS^R495X^ fibrils, at 6 months post-injection (panel A) immunolabelled with amyloid fibril markers LOC and OC (red) and FUS (green). Yellow arrows indicate co-localization between LOC/OC and FUS cytoplasmic inclusions in fibril-injected mice but not in PBS-injected controls (panel B). **C**. Co-localization between LOC (red) and HA-FUS^R495X^ fibrils (green) 3 days post-injection. DAPI (blue) as nuclear counterstaining. Scale bars: 10 µm, inset: 2 µm. **D**. Quantification of the percentage of FUS aggregates that are A11-positive in HA-FUS^R495X^ fibril injected mFUS^KO^/hFUS^R521H^ mice 6 months post-injection. N=3 animals. **E**. Dot-blot analysis of FUS protein levels in urea insoluble fractions of PBS and HA-FUS^R495X^ fibril injected mFUS^KO^/hFUS^R521H^ mouse homogenates using FUS antibody. **F**. Quantification of FUS protein levels in urea insoluble fractions shown in E.

**Figure S6:**
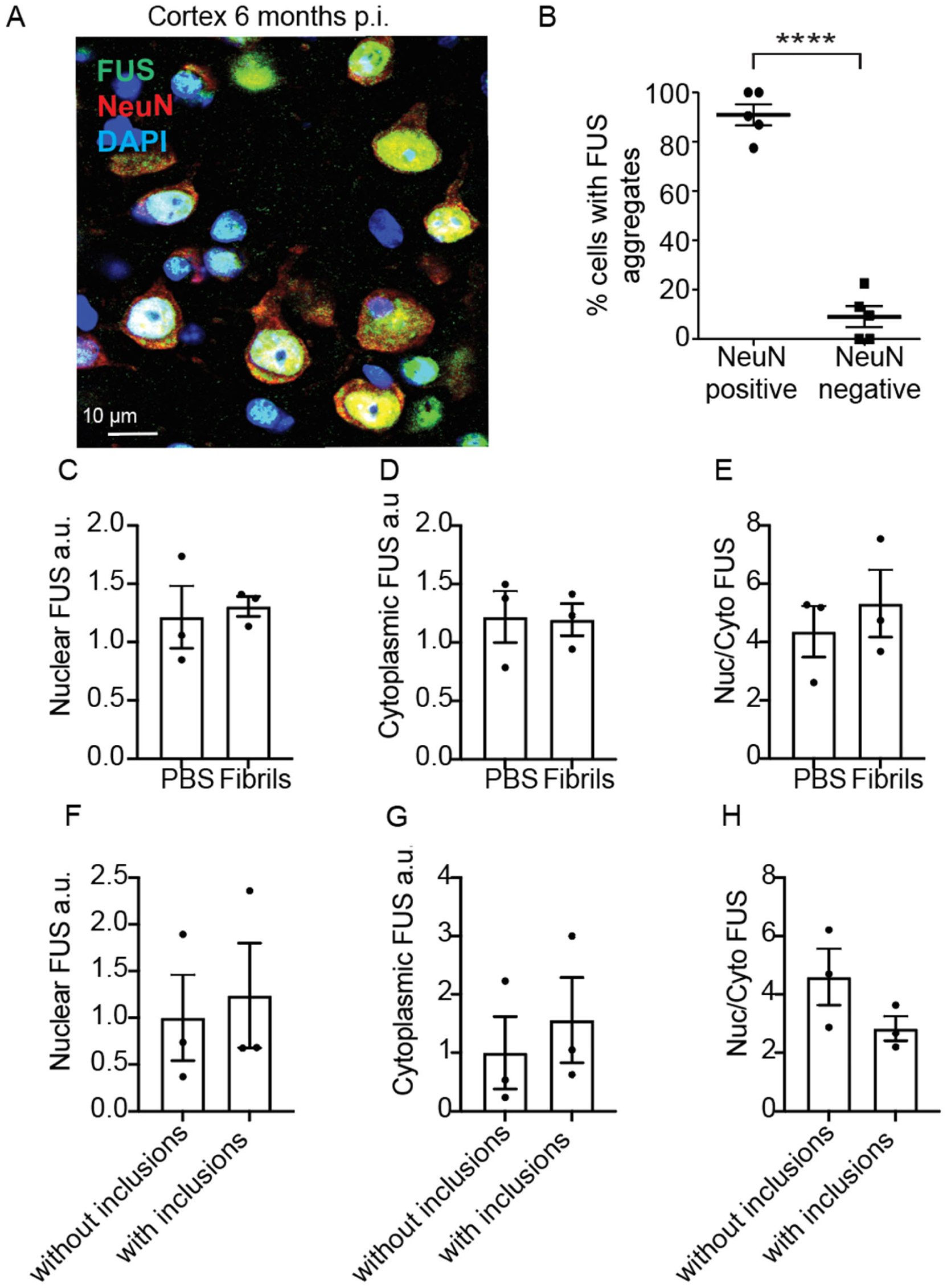
FUS aggregation accumulates predominantly in neurons of fibril-injected mFUS^KO^/hFUS^R521H^ mice. **A.** Immunostaining of mFUS^KO^/hFUS^R521H^ mouse brains injected with HA-FUS^R495X^ fibrils for neuronal marker NeuN (red) and FUS (green) 6 months post-injection. Scale bars: 10 µm. **B.** Quantification of the percentage of NeuN-positive cells and NeuN-negative cells that contain FUS cytoplasmic aggregation. DAPI as nuclear counterstaining. N = 5. **C.** Quantification of the FUS nuclear intensity signal in neurons of both ipsi- and contra-lateral side mice injected either with PBS or with HA-FUS^R495X^ fibrils 6 months post-injection. N = 3 mice per group. **D.** Quantification of the FUS cytoplasmic intensity signal in neurons of both ipsi- and contra-lateral side mice injected either with PBS or with HA-FUS^R495X^ fibrils 6 months post-injection. N = 3 mice per group. **E.** Quantification of the FUS nuclear/cytoplasmic ratio in neurons of both ipsi- and contra-lateral side mice injected either with PBS or with HA-FUS^R495X^ fibrils 6 months post-injection. N = 3 mice per group. **F.** Quantification of nuclear FUS, **G.** cytoplasmic FUS and **H.** FUS nuclear/cytoplasmic ratio in neurons with and without FUS cytoplasmic inclusions of mice injected with HA-FUS^R495X^ fibrils 6 months post-injection. N = 3 mice.

**Figure S7:**
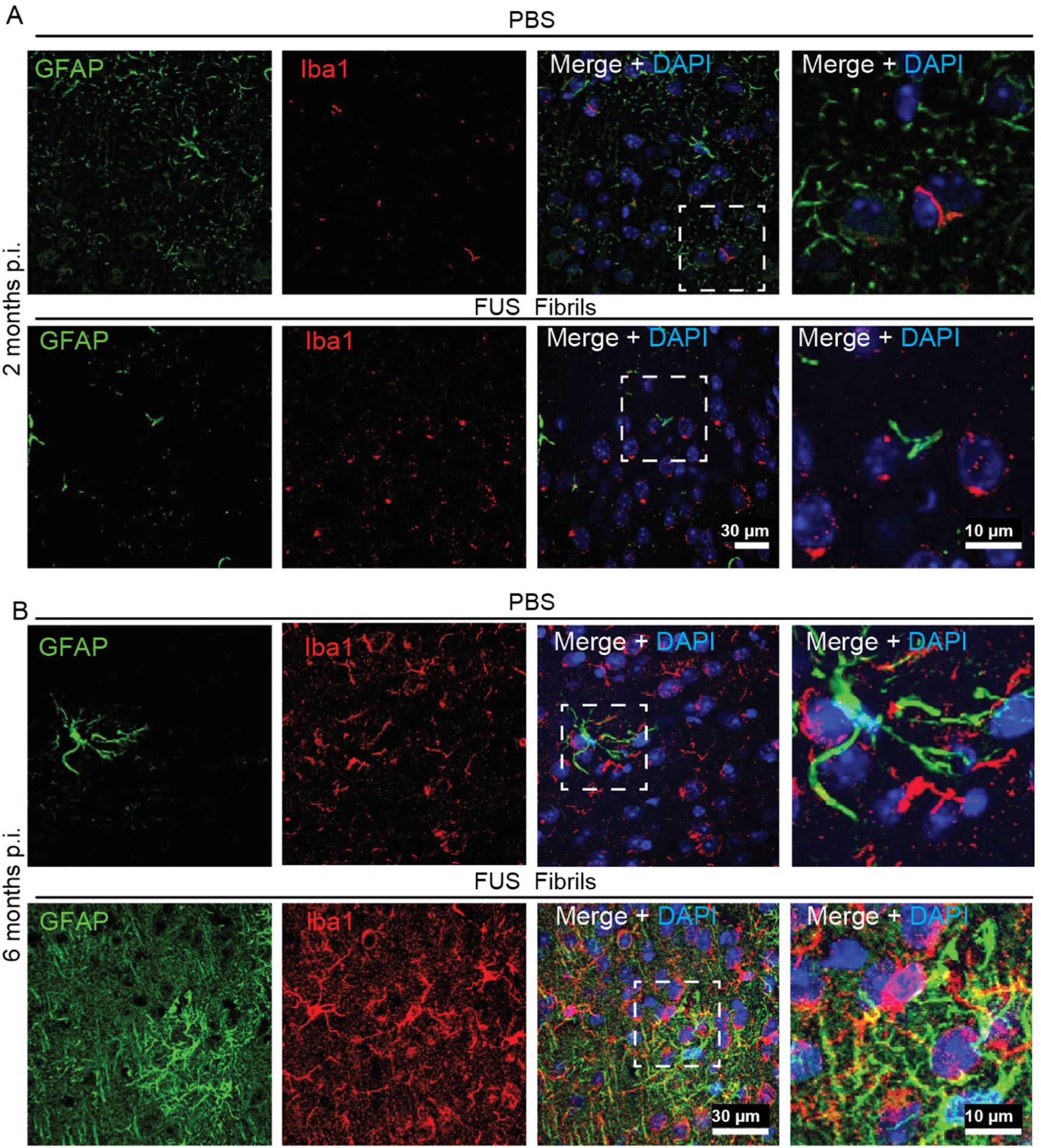
FUS aggregation is accompanied by increased gliosis in fibril-injected mFUS^KO^/hFUS^R521H^ mice. Immunostaining for astrocyte marker GFAP (green) and microglial marker Iba1 (red) of cortex of mFUS^KO^/hFUS^R521H^ mice at 2 (**A**) and 6 months (**B**) post-injection either with PBS or with HA-FUS^R495X^ fibrils. Scale bars: 30 and 10 µm.

**Figure S8:**
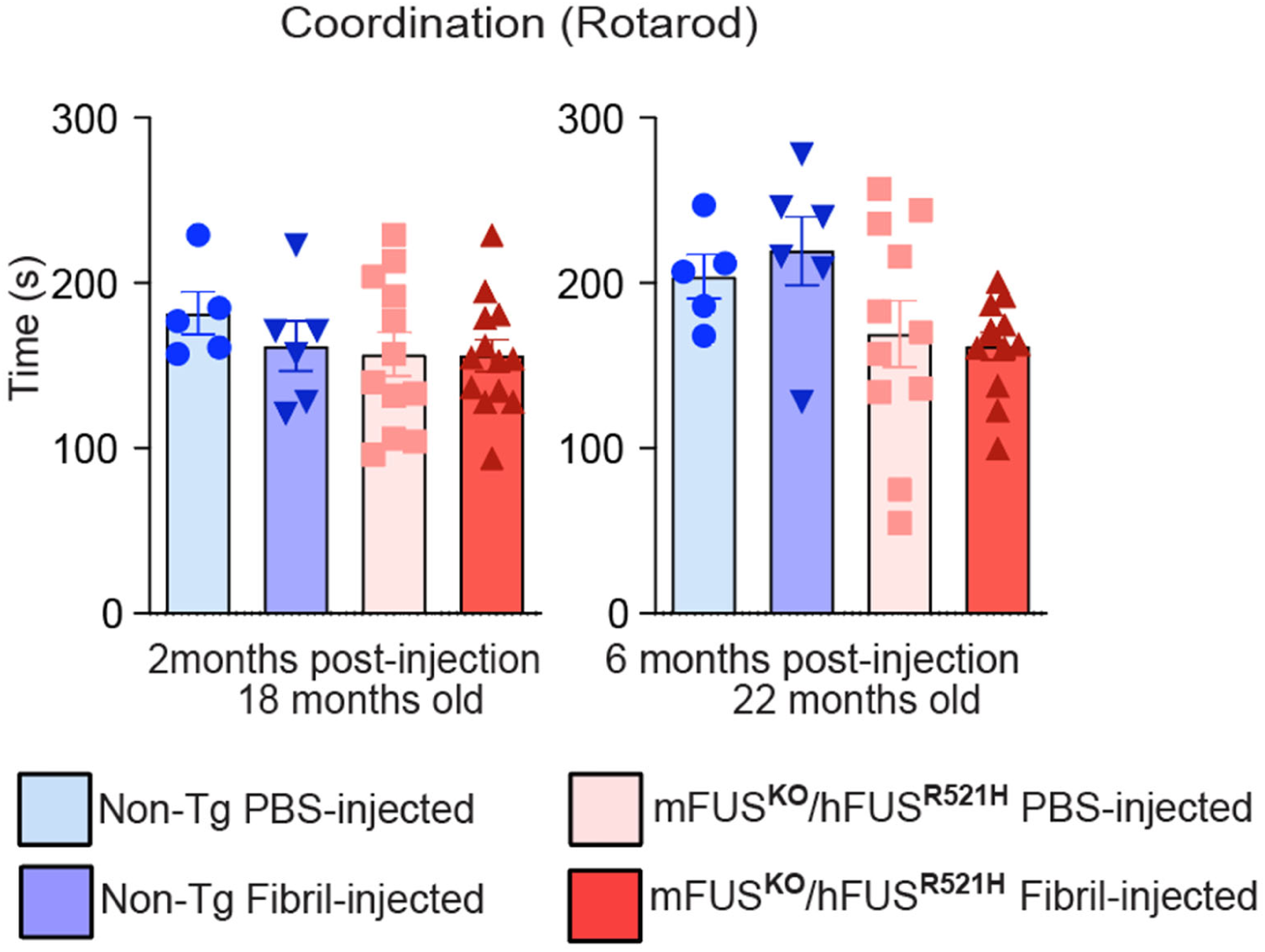
Rotarod performance in non-transgenic and humanized mutant FUS mice injected either with PBS or sonicated FUS fibrils. Rotarod test was performed in 22 months old HA-FUS^R495X^ fibril injected mFUS^KO^/hFUS^R521H^ animals (2 and 6 months post-injection) compared to PBS-injected controls and non-transgenic HA-FUS^R495X^ fibrils or PBS injected controls. N=5–12 animals per group. Data is presented as mean ± SEM.

**Figure S9:**
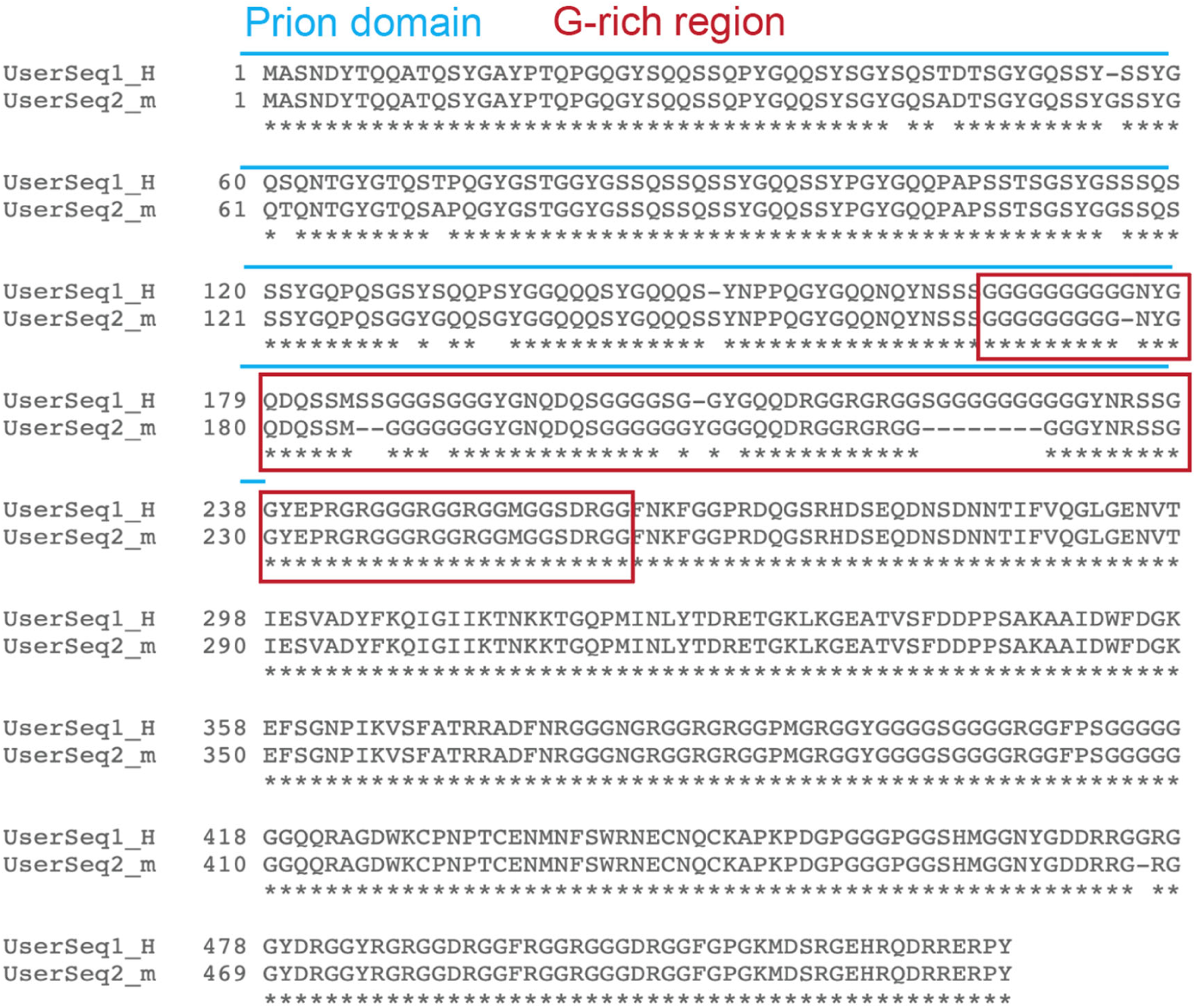
Comparison of human and mouse FUS protein sequences

**Supplementary Table S1.**
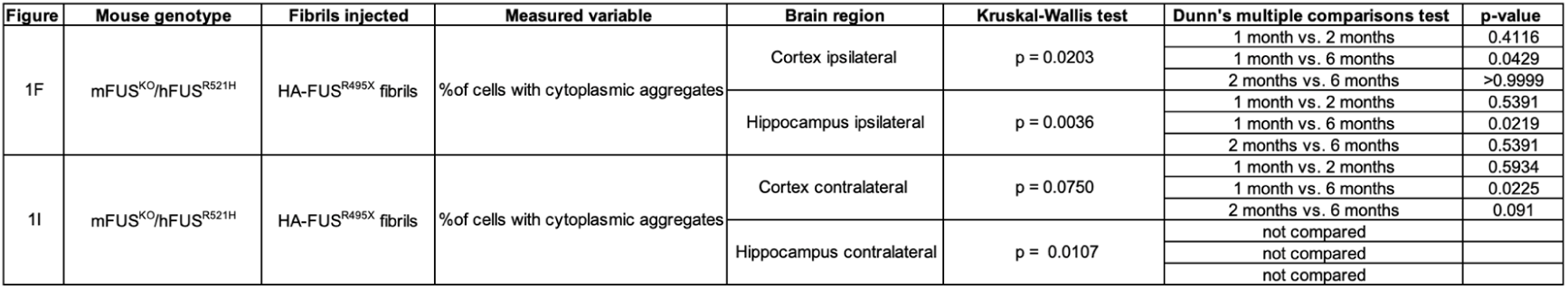
Statistical summary corresponding to Figure 1.

**Supplementary Table S2.**
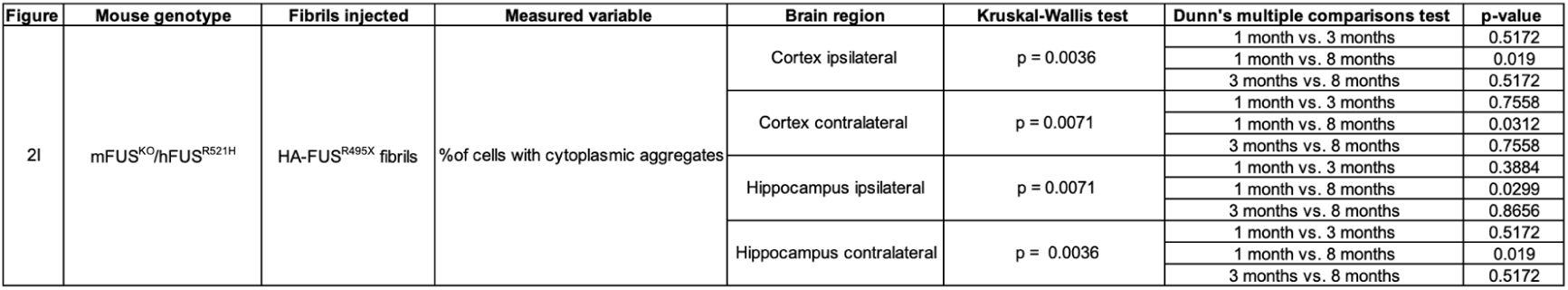
Statistical summary corresponding to Figure 2.

| Figure | Mouse genotype | Fibrils injected | Measured variable | Brain region | Kruskal-Wallis test | Dunn's multiple comparisons test | p-value |
| --- | --- | --- | --- | --- | --- | --- | --- |
| 2I | mFUS<sup>KO</sup>/hFUS<sup>R521H</sup> | HA-FUS<sup>R495X</sup> fibrils | %of cells with cytoplasmic aggregates | Cortex ipsilateral | p = 0.0036 | 1 month vs. 3 months | 0.5172 |
|  |  |  |  |  |  | 1 month vs. 8 months | 0.019 |
|  |  |  |  |  |  | 3 months vs. 8 months | 0.5172 |
|  |  |  |  | Cortex contralateral | p = 0.0071 | 1 month vs. 3 months | 0.7558 |
|  |  |  |  |  |  | 1 month vs. 8 months | 0.0312 |
|  |  |  |  |  |  | 3 months vs. 8 months | 0.7558 |
|  |  |  |  | Hippocampus ipsilateral | p = 0.0071 | 1 month vs. 3 months | 0.3884 |
|  |  |  |  |  |  | 1 month vs. 8 months | 0.0299 |
|  |  |  |  |  |  | 3 months vs. 8 months | 0.8656 |
|  |  |  |  | Hippocampus contralateral | p = 0.0036 | 1 month vs. 3 months | 0.5172 |
|  |  |  |  |  |  | 1 month vs. 8 months | 0.019 |
|  |  |  |  |  |  | 3 months vs. 8 months | 0.5172 |

**Supplementary Table S3.**
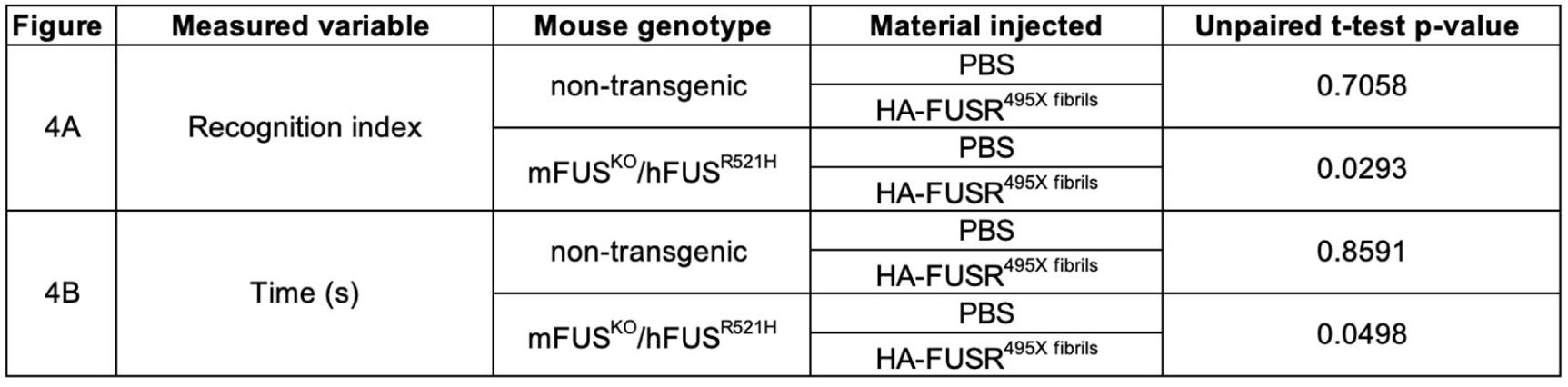
Statistical summary corresponding to Figure 4A and 4B.

| Figure | Measured variable | Mouse genotype | Material injected | Unpaired t-test p-value |
| --- | --- | --- | --- | --- |
| 4A | Recognition index | non-transgenic | PBS | 0.7058 |
|  |  |  | HA-FUS<sup>R495X</sup> fibrils |  |
|  |  | mFUS<sup>KO</sup>/hFUS<sup>R521H</sup> | PBS | 0.0293 |
|  |  |  | HA-FUS<sup>R495X</sup> fibrils |  |
| 4B | Time (s) | non-transgenic | PBS | 0.8591 |
|  |  |  | HA-FUS<sup>R495X</sup> fibrils |  |
|  |  | mFUS<sup>KO</sup>/hFUS<sup>R521H</sup> | PBS | 0.0498 |
|  |  |  | HA-FUS<sup>R495X</sup> fibrils |  |

**Supplementary Table S4.**
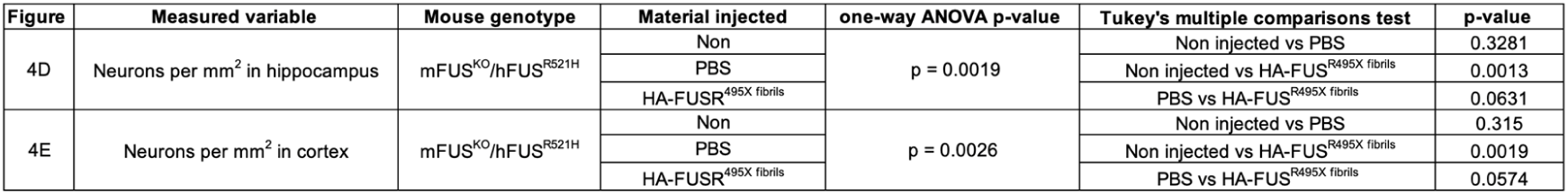
Statistical summary corresponding to Figure 4D and 4E.

**Supplementary Table S5.** Statistical summary corresponding to Supplementary S5.

| Figure | Measured variable | Mouse genotype | Material injected | Compared groups | Unpaired t-test p-value |
| --- | --- | --- | --- | --- | --- |
| S5F | FUS intensity (a.u.) in urea fraction | mFUS<sup>KO</sup>/hFUS<sup>R521H</sup> | HA-FUS<sup>R495X</sup> fibrils | PBS | 0.0024 |
|  |  |  |  | Fibril injected |  |

**Supplementary Table S6.** Statistical summary corresponding to Supplementary S6.

| Figure | Measured variable | Mouse genotype | Material injected | Compared groups | Unpaired t-test p-value |
| --- | --- | --- | --- | --- | --- |
| S6B | % of cells with FUS aggregates | mFUS<sup>KO</sup>/hFUS<sup>R521H</sup> | HA-FUS<sup>R495X</sup> fibrils | NeuN - | <0.0001 |
|  |  |  |  | NeuN + |  |
| S6C | FUS nuclear intensity (a.u.) | mFUS<sup>KO</sup>/hFUS<sup>R521H</sup> | HA-FUS<sup>R495X</sup> fibrils | PBS | 0.7603 |
|  |  |  |  | Fibril injected |  |
| S6D | FUS cytoplasmic intensity (a.u.) | mFUS<sup>KO</sup>/hFUS<sup>R521H</sup> | HA-FUS<sup>R495X</sup> fibrils | PBS | 0.9303 |
|  |  |  |  | Fibril injected |  |
| S6E | FUS nuclear/cytoplasmic ratio | mFUS<sup>KO</sup>/hFUS<sup>R521H</sup> | HA-FUS<sup>R495X</sup> fibrils | PBS | 0.5428 |
|  |  |  |  | Fibril injected |  |
| S6F | FUS nuclear intensity (a.u.) | mFUS<sup>KO</sup>/hFUS<sup>R521H</sup> | HA-FUS<sup>R495X</sup> fibrils | without inclusions | 0.7592 |
|  |  |  |  | with inclusions |  |
| S6G | FUS cytoplasmic intensity (a.u.) | mFUS<sup>KO</sup>/hFUS<sup>R521H</sup> | HA-FUS<sup>R495X</sup> fibrils | without inclusions | 0.5919 |
|  |  |  |  | with inclusions |  |
| S6H | FUS nuclear/cytoplasmic ratio | mFUS<sup>KO</sup>/hFUS<sup>R521H</sup> | HA-FUS<sup>R495X</sup> fibrils | without inclusions | 0.1695 |
|  |  |  |  | with inclusions |  |

**Supplementary Table S7.** Statistical summary corresponding to Supplementary S8.

| Figure | Measured variable | Mouse genotype | Material injected | Unpaired t-test p-value |
| --- | --- | --- | --- | --- |
| S8 | Time (s) | non-transgenic | PBS | 0.5656 |
|  |  |  | HA-FUSR<sup>495X</sup> fibrils |  |
|  |  | mFUS<sup>KO</sup>/hFUS<sup>R521H</sup> | PBS | 0.7252 |
|  |  |  | HA-FUSR<sup>495X</sup> fibrils |  |

